# The Ribosome Profiling landscape of yeast reveals a high diversity in pervasive translation

**DOI:** 10.1101/2023.03.16.532990

**Authors:** Chris Papadopoulos, Hugo Arbes, Nicolas Chevrollier, Sandra Blanchet, David Cornu, Paul Roginski, Camille Rabier, Safiya Atia, Olivier Lespinet, Olivier Namy, Anne Lopes

## Abstract

Pervasive translation is a widespread phenomenon that plays an important role in de novo gene birth; however, its underlying mechanisms remain unclear. Based on multiple Ribosome Profiling (Ribo-Seq) datasets, we investigated the RiboSeq landscape of coding and noncoding regions of yeast. Therefore, we developed a representation framework which allows the visual representation and rational classification of the entire diversity of Ribo-Seq signals that could be observed in yeast. We show that if coding regions are restricted to specific areas of the Ribo-Seq landscape, noncoding regions are associated with a wide diversity of translation signals and, conversely, populate the entire yeast Ribo-Seq landscape. Specifically, we reveal that noncoding regions are associated with canonical translation signals, but also with non-canonical ones absent from coding regions, and which appear to be a hallmark of pervasive translation. Notably, we report thousands of translated noncoding ORFs among which, 251 led to detectable products with Mass Spectrometry while being characterized by a wide range of translation specificities. Overall, we show that pervasive translation is not random with noncoding ORF translation signals being consistent across Ribo-Seq experiments. Finally, we show that the translation signal of noncoding ORFs is not explained by features related to the emergence of function, but rather determined by the translation start codon and the codon distribution in their two alternative frames. Overall, our results enable us to propose a topology of the pervasive Ribo-Seq landscape of a species, and open the way to future comparative analyses of this translation landscape under different conditions.

## Introduction

All organisms undergo molecular innovation to adapt to their environment. Typically, molecular innovation involves the creation of novel products or the formation of novel interactions between already existing products (i.e., genes, proteins). The former case implies the existence of a “reservoir” of freely and/or fast-evolving sequences, which allows for the sampling of extensive sequence spaces that could not be reached under natural selection. In theory, to ensure the efficiency of the functional processes that occur in the cell, this reservoir is expected to be locked (i.e., not expressed), to avoid unselected products that could interfere with the established ones. Nevertheless, from an evolutionary perspective, the pervasivity of biological processes (i.e., transcription, translation…) enables the expression of small fractions of novel sequence spaces, thereby exposing them to selection, and allowing from time to time, the passage from this reservoir to the world of established, regulated, and selected products.

In fact, noncoding regions can be seen as a reservoir of unselected sequences, hosting thousands of small Open Reading Frames (ORFs) that could give rise to novel products if translated ^1–6^. Precisely, OMICS technologies have provided a huge amount of data revealing the “omnipresence” of biological noise which has turned out to result from the pervasivity of biological processes. As a matter of fact, noncoding regions have been shown to be pervasively transcribed and translated, exposing non-genic sequences to selection ^4,7–25^. Furthermore, hundreds of novel peptides or microproteins resulting from presumed noncoding regions have been detected with proteomics, further supporting the pervasivity of translation ^19,26–32^. Finally, many studies report examples of de novo gene birth from noncoding regions in eukaryotic species ^24,33–58^. These de novo genes exhibit clear regulation patterns, have been shown to be subject to negative selection and a function has been reported for some of them, confirming that they could undoubtedly be associated with the coding world ^34,35,43,57,59,60^. If different models of de novo gene emergence have been proposed so far, most of them share the hypothesis of an early stage as a protogene or small peptide that results from the translation of noncoding regions ^10,20,21,44,47,50,54^. These models consequently attribute an important role to pervasive translation in de novo gene birth, constituting somehow the last step for a noncoding ORF to reach the protein state, prerequisite, though not sufficient, to enter the coding world. All the aforementioned studies therefore (i) reveal that the passage from the noncoding to the coding world is much more frequent than previously thought, and (ii) place the noncoding genome, but also pervasivity at the center of the emergence of genetic novelty enabling, from time to time, the passage from unselected products to functional and regulated ones.

Each RNA molecule contains three potential reading frames, each one hosting distinct but overlapping ORFs which encode different amino acid sequences. In coding regions, whose translation is strongly regulated to ensure the production of the functional protein, the translation is unbalanced toward the frame hosting the coding sequence (CDS) so that the translation outcome is most of the time the coding product. This leads to a remarkable triplet periodicity of Ribosome Profiling (Ribo-Seq) reads, the two CDS alternative frames being devoid of reads. As such, the canonical translation patterns of regulated translation are easily detectable by Ribo-Seq analysis tools. However, in noncoding regions undergoing pervasive, and therefore nonregulated translation, the nature and diversity of translation signals remain unknown. Yet, pervasive translation is usually studied with methods dedicated to coding regions, based themselves on rules established from coding regions. Specifically, the focus is put on canonical translation signals that resemble those of coding sequences undergoing regular translation. Nonetheless, pervasive translation events that participate in the generation of de novo products might be found in the discarded non-canonical translation signals which, therefore, deserve to be considered. In order to explore the extent to which pervasive translation can give rise to novel protein products and the mechanisms governing their production, one needs to delineate the boundaries of pervasive translation, and characterize the diversity of pervasive translation signals, including both canonical and noncanonical ones. To do so, we devised a representation framework that maps all the ORFs lying in ribosome-associated RNAs on a 2D plan according to their fractions of reads in their own and alternative frames, respectively, without any *a priori* on their translation outcome. Detecting pervasively translated ORFs events is very challenging due to their short size and typically low expression levels. Furthermore, these ORFs are not expected to be expressed constitutively, but instead, are likely to be translated in only a few experiments. Therefore, to increase the ribosome profiling signal and cover the diversity of pervasive translation events, we took advantage of the accumulation of Ribo-Seq data on yeast, and collected 101 Ribo-Seq datasets. We then characterized the translation activity of the whole genome of *S. cerevisiae* including both coding regions (i.e., CDSs and their alternative frames) and noncoding ones. The translation signals of coding regions not only enabled us to establish a reference for studying the characteristics of pervasive translation, but also allowed us to identify non-canonical translation events in coding regions. We then performed Mass Spectrometry (MS) to investigate whether the different types of translation signals could be associated with a peptide. Finally, we investigated the features that could play a role in the final translation outcome of a noncoding ORF. Therefore, we characterized the properties of noncoding ORFs with respect to their translation status to better understand the rules, if any, that dictate their translation outcome and that finally, would underlie pervasive translation.

## Results

### Collection of Ribo-Seq experiments

Ribo-Seq provides a genome-wide picture of the RNAs undergoing translation^8^. Briefly, the method consists in sequencing, after RNA digestion, the fragments (i.e., reads) that were protected by the translating ribosomes, thereby identifying the RNA regions that were bound by ribosomes. High quality Ribo-Seq experiments can detect the translation signal at the single nucleotide resolution, allowing for the precise detection of the translated codons and, therefore, of the RNA frame that was translated. In coding regions, since the coding frame is known, the translated frame is straightforward to detect even when dealing with Ribo-Seq experiments of intermediate or even poor quality. In contrast, in noncoding regions, identifying the translated frame in noncoding regions requires high-quality Ribo-Seq data since there is no prior knowledge on the frame that is translated. Furthermore, noncoding ORFs are usually expressed at low level and not expected to be translated systematically. Finally, we aim to delineate the pervasive translation that can be expected under standard conditions and not to be biased by experimental conditions or mutants that could induce translation deregulation. Therefore, to enhance our ability to detect low Ribo-Seq signals, and capture both occasional and constitutive translation events that may occur under standard conditions, we assembled a significant collection of 101 publicly available Ribo-Seq datasets performed in standard conditions on the wild-type S288C or BY4741 strains of *Saccharomyces cerevisiae*, from which we were able to extract 58 high quality datasets (see Methods).

### Representation framework

We aim to inventory the diversity of translation signals observed in pervasive translation and compare them to those of yeast established genes. To do so, we divided the yeast genome into two genomic categories: (i) the coding regions, including the annotated CDSs and all the noncoding ORFs hosted in the CDSs’ alternative frames (96873 aORFs for alternative ORFs), and (ii), the noncoding regions, which consist in a set of 78989 ORFs located in the inter-CDS regions (iORFs for inter-CDS ORFs) that do not overlap any annotated feature of *S. cerevisiae* genome (Methods – Table S1). aORFs and iORFs have both a minimal size of 60 nucleotides which is a reasonable size for generating a microprotein that can be fixed and established as novel protein. In order to also detect non-canonical translation initiation events, noncoding ORFs do not necessarily start with ATG codons and are defined from STOP to STOP. It is worth noting that our pipeline can handle any ORF type, regardless of their boundaries, as long as they are specified in a GFF file. The reads extracted from the 58 Ribo-Seq runs were then pooled and mapped to the CDSs, aORFs, and iORFs with our pipeline ORFribo (see Methods and Figure S1). Beyond detecting ORFs translated with high specificity, we aim to capture the full diversity of pervasive translation events and, therefore, probe the entire range of Ribo-Seq signals in ribosome-bound RNAs. To do so, we retained all the ORFs associated with at least 50 reads, regardless of the frame, and whose read codon coverage was above 30%. This allowed us to discard cases where reads accumulate on very few codons that may result from repartition bias due to experimental artifacts.

For each ORF, we calculated its fraction of in-frame reads (reads mapping to the frame of the ORF, named by convention frame 0 (F0)) and that of its out-of-frame reads mapping to its +1 and +2 frames (F1 and F2 respectively) (Figure 1A). The higher its fraction of in-frame reads, the higher its specificity of translation. Translated CDSs are expected to display canonical translation signals with a good in-frame read triplet periodicity, where each codon is associated with a high fraction of F0 reads. On the opposite, their aORFs are expected to exhibit shifted periodicity with a majority of reads in F1 or F2 that in fact reflect the translation of the overlapping CDS. However, the translation patterns associated with RNAs undergoing pervasive translation remain unknown. We, therefore, devised a representation framework that provides the Ribo-Seq landscape of a set of ORFs of interest. In this 2D Ribo-Seq landscape, each ORF is positioned according to its fractions of reads in the F0 and F1 frames (Figure 1B). The x-axis of the Ribo-Seq landscape is divided into three equivalent regions to distinguish (i) high-FIR ORFs (i.e. high Fraction of In-frame Reads) whose reads are mostly in-frame, reflecting ORFs translated with high specificity, (ii) low-FIR ORFs whose reads are mostly out-of-frame, which, in theory, reflect untranslated ORFs overlapping translated ones (they may nevertheless be associated with a few in-frame reads due to misassignment of the translated frame), and (iii), intermediate-FIR ORFs with an intermediate translation specificity that display enrichment in in-frame reads compared to what could be expected if the three frames were equivalently translated (F0 fraction ≥ 0.33), while not specifically translated. As such, the x-axis reflects the specificity of translation of a given ORF, while the y-axis enables one to estimate the ribosome occupancy in its two alternative frames.

**Figure 1:**
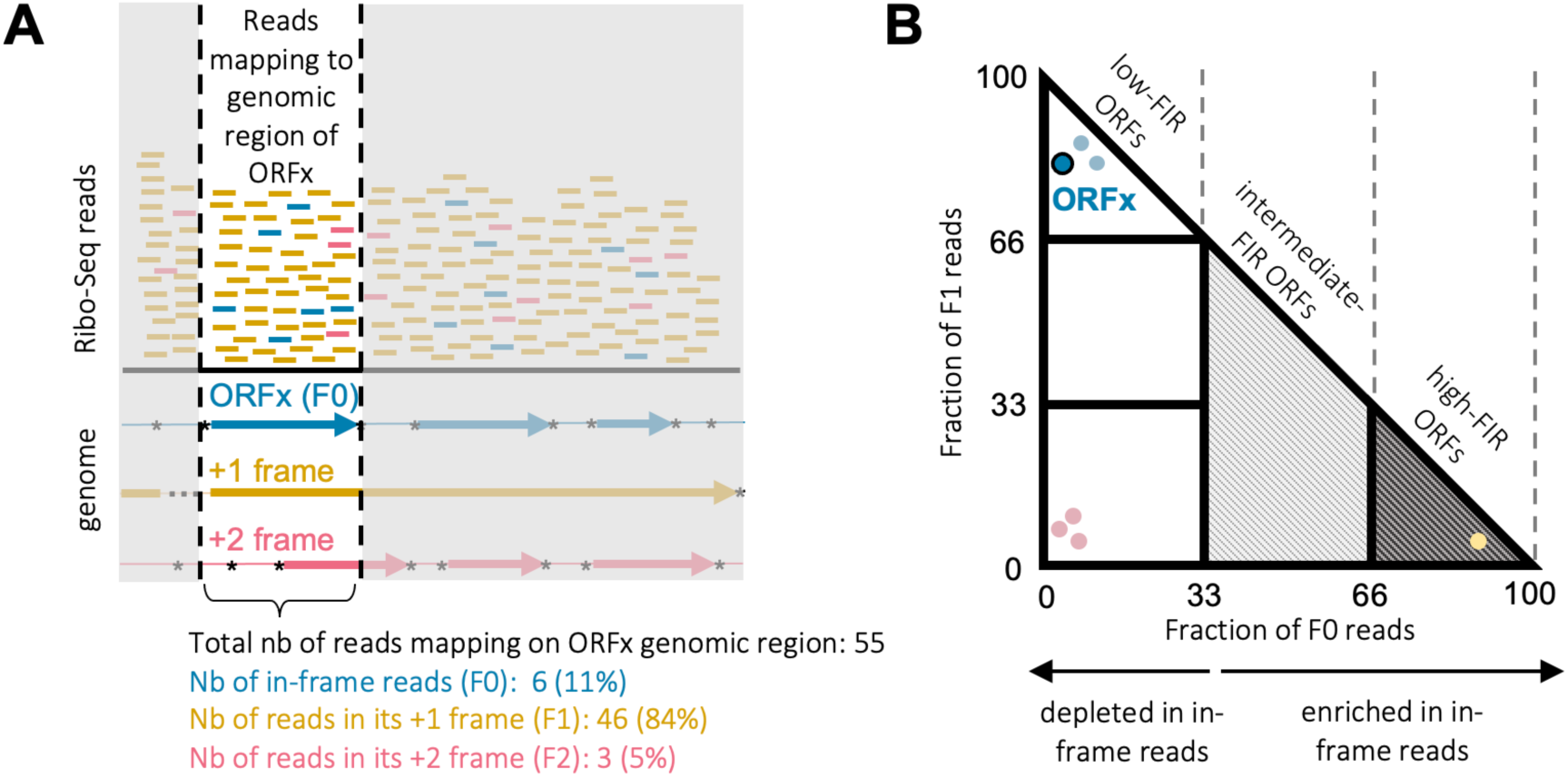
Construction of the Ribo-Seq landscape. **(A)** Example of calculation of the fractions of F0, F1, F2 reads of ORF X. Top: Each Ribo-Seq read according to its genomic coordinates can be assigned to a unique nucleotide, thereby indicating the codon and thus the frame that was translated by the ribosome. They are mapped to the genome and colored according to their corresponding frame. Bottom: Genome annotation with STOPs indicated by stars, and ORFs with at least 60 nucleotides represented with arrows. Reads mapping to the genomic region of ORF X, no matter the frame, are surrounded by vertical lines. 6 of them are in-frame, mapping on the reading frame of X and counted as F0 reads. 46 of them are out-of-frame, mapping on the +1 frame of X and counted as F1 reads. The remaining 3 reads are also out-of-frame, mapping to the +2 frame of X, and counted as F2 reads. **(B)** Each ORF is represented on the Ribo-Seq landscape according to its fractions of reads in its F0 and F1 frames (the fraction of reads in its +2 frame (F2) can be directly deduced from those of F0 and F1). ORF X is surrounded in black. Theoretically, random translation is expected to lead to reads evenly distributed across the three frames leading approximately to F0 = F1 = F2 = 33%. We considered as enriched in a given frame, all ORFs associated with a fraction of in-frame reads higher than 33%. ORF translation status (low-, intermediate- and high-FIR) are defined according to their fractions of reads in their F0 frame (i.e., fraction of reads in the reading frame of the considered ORF) and reflect different levels of translation specificities.

### Ribo-Seq landscape of coding regions

As expected, most of the read-associated CDS (5403 - 93%) fall within the high-FIR region of the Ribo-Seq landscape (F0 ≥ 0.66), highlighting their highly specific translation (Figure 2A, Table S1). This is evidenced by their metagene profile, which displays a strong triplet periodicity with reads accumulating in their F0 frame (Figure 2B). Conversely, 98% (76449) of aORFs associated with at least 50 reads exhibit high fractions of out-of-frame reads (fraction of F0 reads < 0.33) and locate at the two left extremities of the Ribo-Seq landscape. The out-of-frame reads of aORFs probably do not reflect a translation outcome, but instead mirror the translation of the overlapping CDSs since the profiles of overlapping ORFs are inherently correlated. This is supported by their metagene profiles where reads accumulate in the F1 and F2 frames (Figure 2B). Overall, the three extremities of the Ribo-Seq landscape highlight RNA regions associated with an unbalanced ribosome occupancy between the three RNA frames in favor of that of the CDS. We will refer to these regions as regions of unbalanced ribosome occupancy.

**Figure 2:**
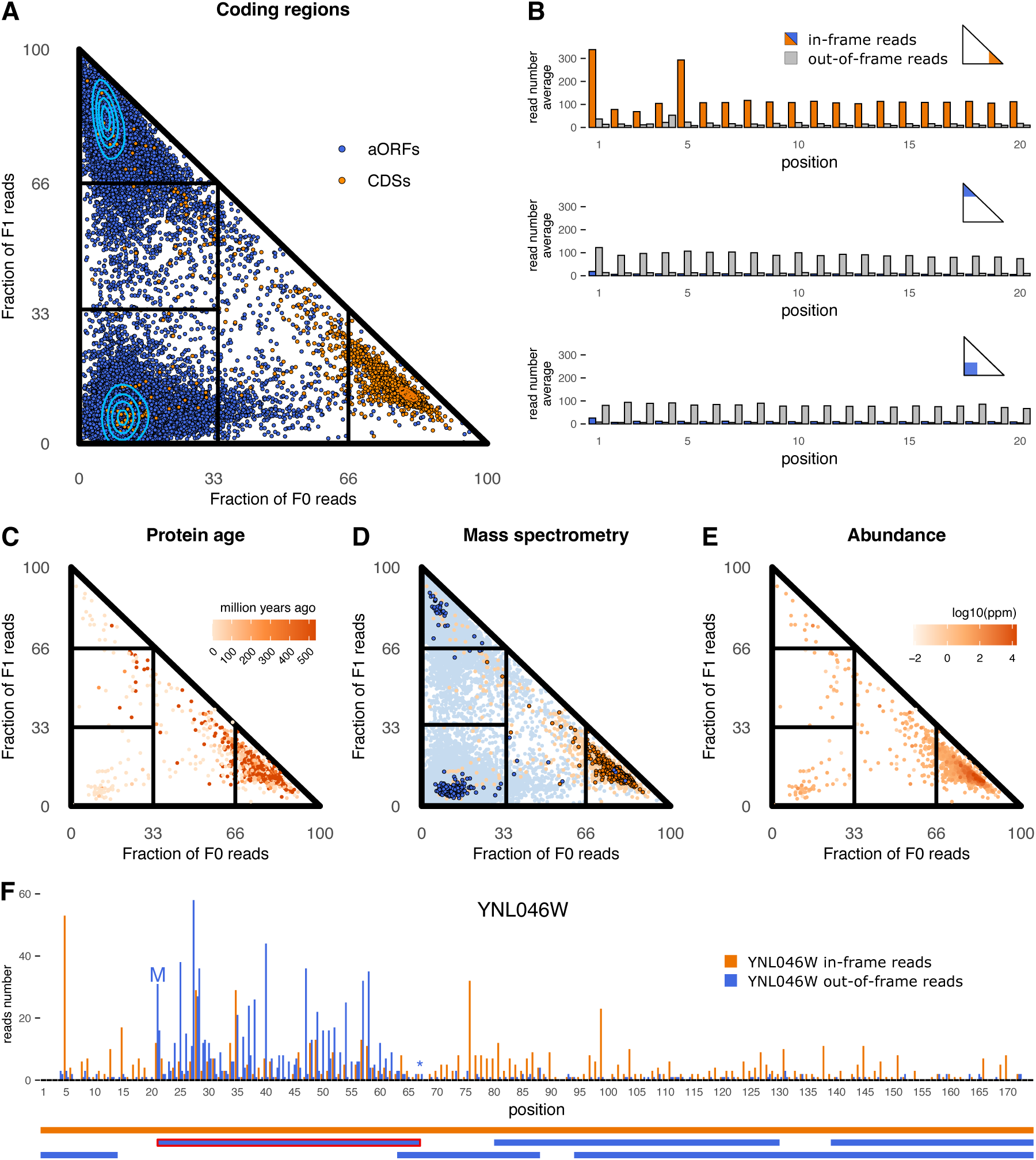
Ribo-Seq landscape of coding regions. **(A)** Ribo-Seq landscape of the coding regions’ ORFs, including CDSs and aORFs, colored orange and blue, respectively. Each ORF with at least 50 reads and 30% of codon coverage is represented as a dot on the Ribo-Seq landscape according to its fractions of F0 reads (x-axis) and F1 reads (y-axis). **(B)** Metagenes of the three extremities of the Ribo-Seq landscape. Each metagene is calculated for the first 20 codons from all the ORFs associated with a given Ribo-Seq area (see Methods) and represents the average number of reads per nucleotide over all the ORFs of the metagene. The corresponding Ribo-Seq landscape areas are colored on the schematic Ribo-Seq landscapes of each plot. Nucleotide positions that are covered by in-frame reads relative to the ORFs of the metagene are colored in orange or blue, while positions covered by out-of-frame reads (F1 and F2) are colored in grey. **(C)** Ribo-Seq landscape of the CDSs colored with respect to their evolutionary age (in Mya) as estimated by phylostratigraphy (see Methods). **(D)** Ribo-Seq landscape of the aORFs and CDSs observed in our MS data. Same representation as panel A, but aORFs and CDSs for which we detected a peptide product are colored in dark blue or dark orange, respectively. **(E)** Ribo-Seq landscape of the CDSs colored according to their protein abundance in ppm as provided by PaxDb ^61^. **(F)** Top: Metagene of the CDS YNL046W representing the number of reads of each nucleotide summed over the 58 Ribo-Seq datasets. Positions that are covered by reads in frame with the CDS YNL046W are colored in orange, while those covered by reads in its +1 and +2 frames are colored in blue. Bottom: Colored rectangles represent the CDS (orange) and its aORFs (blue). The start and stop codons of aORF chrXIV:542365-542502 (surrounded in red) are indicated by a “M” and a star, respectively.

Interestingly, a small fraction of CDSs (165 cases ∼3%) fall within the intermediate- and low-FIR regions of the Ribo-Seq landscape (F0 < 0.66). 64% (107) of them are specific to the Saccharomyces *sensu stricto* (Figure 2C) with 50% (53) overlapping on the same strand another CDS that is older and specifically translated. This suggests young or emerging ORFs that have not yet acquired the regulatory elements necessary for efficient expression, being therefore associated with ribosomes that share their occupancy with the frame of the older overlapping CDS. However, 65% (108) of the CDSs associated with high fractions of out-of-frame reads do not overlap another CDS. We carefully analyzed their individual metagene profiles and present an illustrative example where the ribosome footprints are shared between a CDS and one of its aORFs (Figure 2F). Interestingly, if in-frame reads cover the whole sequence of the CDS YNL046W, reads in the +1 frame accumulate between the codons 21 and 67. This region precisely corresponds to the genomic coordinates of the aORF chrXIV:542365-542502 which is enriched in in-frame reads (fraction of F0 reads: 57%). The first peak of in-frame reads observed for the aORF lies at an ATG codon and probably indicates the translation initiation. The fact that the high density of out-of-frame reads of the gene YNL046W overall correlates with the borders delineated by the first ATG and the STOP codon of the aORF, and that no comparable signal is observed along the remaining regions of the CDS, strongly supports that this signal does not reflect noise but rather the translation of this aORF. In fact, a significant portion (65/165) of the low- or intermediate-FIR CDSs overlap aORFs associated with intermediate or high fractions of in-frame reads, which could indicate an alternative translation outcome, although this remains to be fully demonstrated. However, it is unclear whether these cases reflect distinct RNA molecules that are independently translated, a competition occurring at the translation level between overlapping ORFs, or independent translation events from the same RNA molecule. Nonetheless, our results suggest that these ORFs, despite their non-canonical translation signals, might be associated with a translation outcome and warrant further investigation.

### Detection of translated products with Mass Spectrometry

In order to verify whether ORFs (including CDSs and aORFs) with intermediate or low fractions of in-frame reads could be associated with a protein product, we undertook MS experiments in standard conditions and conditions where the proteasome is inhibited (see Methods). Indeed, aORFs are short, may encode unstable peptides and if translated, the abundance of their resulting peptides is expected to be very low, thereby rendering their detection very challenging. Inhibiting the proteasome should, therefore, increase our chances of detecting unstable and/or short-lived peptides that are normally rapidly degraded by the proteasome. We detected at least two non-redundant peptides for 3716 high-FIR CDSs and 10 CDSs associated with intermediate and low-FIR regions of the Ribo-Seq landscape, supporting that the latter may occasionally be related to translation events leading to detectable products (Figure 2D). We then extracted CDS protein abundances from the PaxDb database ^61^, a curated meta-resource that integrates multiple mass spectrometry datasets for which abundances were reprocessed, unified, and scored (Figure 2E). Most of the low- and intermediate-FIR CDSs (128 - 78%) were detected in the MS data of PaxDb. Although they display lower abundances than high-FIR CDS (one sided Wilcoxon test P = 2.67 × 10^−32^), these results strengthen the hypothesis that CDSs translated with intermediate or low specificity, may still result in a protein product. Finally, when pooling the two conditions (wild-type and inhibited proteasome), we detected at least one peptide for 217 aORFs (Figure 2D, Table S2). Most of these aORFs (211 - 97%) are located in the low-FIR area of the Ribo-Seq landscape, again supporting that this region can occasionally be associated with translation outcomes. Interestingly, these aORFs are associated with Ribo-Seq reads more frequently than other aORFs, display higher numbers of F0 reads and overlap CDSs also characterized by higher numbers of F0 reads than other CDSs (Mann–Whitney U-test (one-sided), P = 2.2 × 10^−14^, 7.3 × 10^−25^ and 1.16 × 10^−12^, respectively) (Figure S2-4). This suggests (i) that these aORFs, benefiting from the higher expression rates of their overlapping CDS, are consequently also translated at higher rates, probably explaining the fact that we were able to detect their resulting peptides, and (ii) that regions with high expression activity are more likely to produce through pervasive translation, a peptide in sufficient quantity to be detected in MS experiments. If it is tempting to speculate that regions of high expression activity are more likely to generate novel coding products, the relationship between the quantity of a peptide and its capacity to ensure a biological role and finally to be fixed is to be further investigated. Moreover, although we probably underestimate the number of peptides resulting from aORF translation, this result does not mean either that all the ORFs lying in the low-FIR region are translated. Indeed, their metagenes strongly suggest a non-translational outcome for the majority of them as observed for most aORFs of the gene YNL046W (Figure 2F). However, our MS data show that a small fraction of them is translated in enough quantities to be detected at the peptide level.

### Ribo-Seq landscape of noncoding regions

We identified 6788 iORFs with at least 50 reads and a read codon coverage of 30%. Contrarily to coding regions, noncoding regions are associated with a wide diversity of translation signals and populate the entire yeast Ribo-Seq landscape. Specifically, we reveal a continuum in the translation signals that is absent from coding regions and appears to be a hallmark of pervasive translation of noncoding regions (Figure 3A). Higher densities of iORFs are nonetheless observed at the three extremities of the Ribo-Seq landscape. Notably we report 1036 high-FIR ORFs and 2244 low-FIR ORFs located at the two left extremities of the Ribo-Seq landscape. The metagene profile of the high-FIR ORFs is typical of canonical CDSs (Figure 3F), suggesting iORFs whose translation clearly dominates that of their alternative frames, and revealing that pervasive translation can nevertheless be associated with high translation specificity. On the other hand, the metagene profiles of the low-FIR iORFs that locate at the two left extremities of the Ribo-Seq landscape resemble those of aORFs, and rather mirror the translation activity occurring in their alternative frames (Figure 3BD). An important fraction of read-associated iORFs (2226 - 33%) falls within the intermediate-FIR region of the Ribo-Seq landscape (0.33 ≤ F0 < 0.66). Theoretically, random translation is expected to be balanced between the three frames with reads evenly distributed across them (i.e., fractions of reads in F0, F1, F2 ∼ 0.33). Instead, this twilight zone gathers iORFs enriched in in-frame reads, although the ribosomes seem to share their occupancy with the alternative frames. This is well supported by their metagene profile, where in-frame reads prevail over the other frames, though the contrast is less pronounced than that observed for high-FIR iORFs (Figure 3E). These iORFs precisely recall the hundred CDSs observed in the same Ribo-Seq landscape region that were translated with a detectable protein product despite their low translation specificity (Figure 2DEF). The remaining area, which is also poorly populated in the Ribo-Seq landscape of coding regions, corresponds to low-FIR ORFs with intermediate fractions of F1 and F2 reads (0.33 ≤ [F1, F2] < 0.66). In fact, these iORFs mainly correspond to iORFs overlapping intermediate-FIR iORFs (one-proportion z-test (one-sided), P = 3 × 10^−22^)(Figure S5). Their corresponding region is also referred to as the twilight zone. Similarly to intermediate-FIR ORFs, their metagene profile is less contrasted than that of the other low-FIR iORFs (Figure 3C). Furthermore, our MS experiments enabled us to detect a peptide for 34 iORFs considering both standard and inhibited proteasome conditions (Methods, Figure 3A and Table S2). The detected peptides correspond to iORFs with different translation specificities, including high-, intermediate- but also low-FIR ORFs. This, again, reinforces that the different Ribo-Seq landscape areas including those associated with translation signals of intermediate or low specificities (i.e., non-canonical) can occasionally be associated with a translation product.

**Figure 3:**
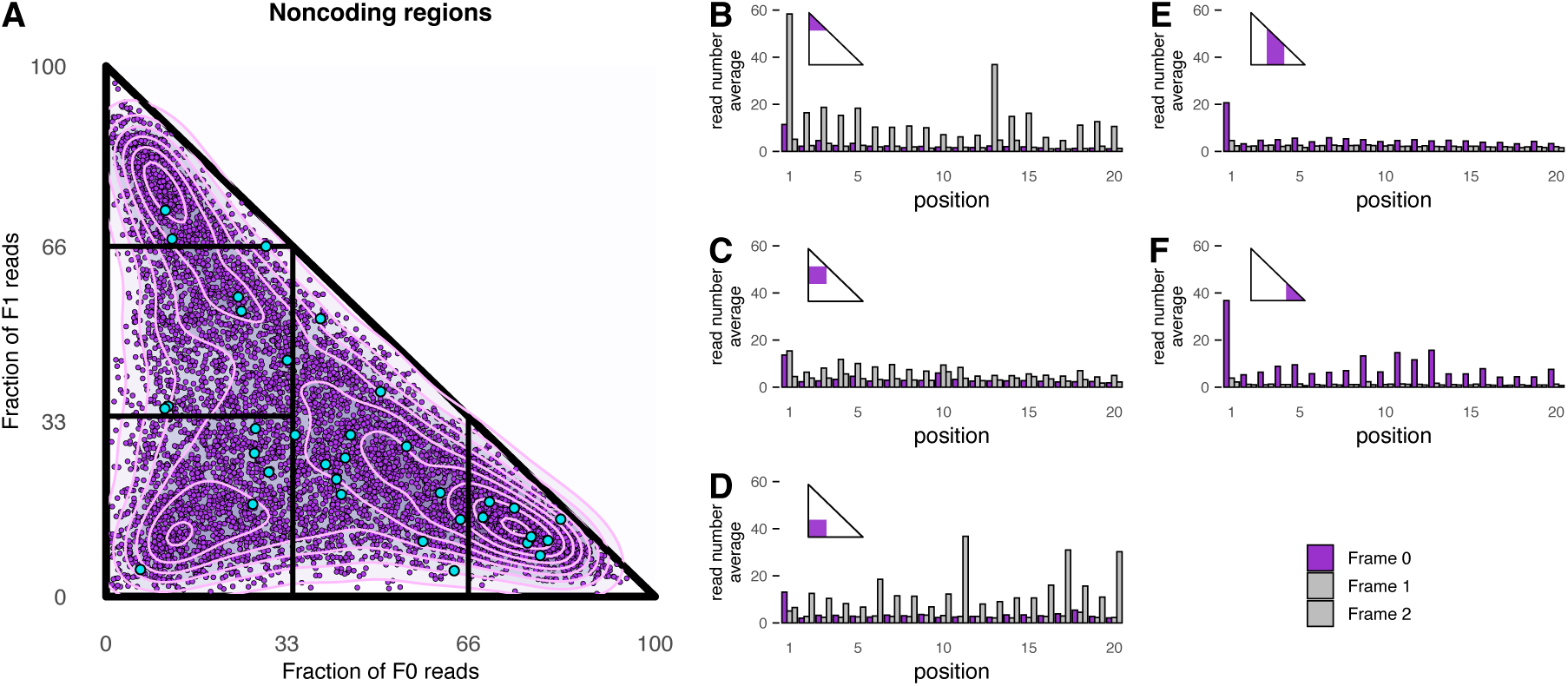
Translational landscape of noncoding regions. **(A)** Ribo-Seq landscape of the iORFs. Each iORF with more than 50 reads and 30% od read codon coverage is represented as a purple dot on the Ribo-Seq landscape according to its fractions of F0 reads (x-axis) and F1 reads (y-axis). Cyan dots correspond to iORFs for which at least one peptide has been detected with MS. **(B-F)** Metagenes of the five regions of the Ribo-Seq landscape representing the average number of Ribo-Seq reads of each nucleotide for the first 20 codons. Metagenes are calculated as in Figure 2 and the Ribo-Seq landscape regions of the ORFs used for the calculation of each metagene are colored in purple in the corresponding schematic Ribo-Seq landscapes. Nucleotide positions that are covered by in-frame reads (F0) for the considered iORFs are represented in purple while those covering reads in the +1 and +2 frames are colored in grey.

### Robustness of the translation signals across Ribo-Seq samples

We investigated the contribution of each dataset to the global Ribo-Seq landscape. Figure 4A shows the distribution of the fractions of the ORFs (CDSs, aORFs or iORFs) of the global Ribo-Seq landscape that were detected in each individual sample. For each sample, an ORF was retained if it was associated with at least 10 reads regardless of the frame (Figure S6). Again, beyond detecting translated ORFs, we aim to compile an inventory of all the ORFs present in ribosome-bound RNAs. Furthermore, while a threshold of 50 reads per sample may be too strict, we believe that 10 reads could be a lower limit for computing fractions of in-frame reads. Most datasets display high coverage for the CDSs present in the global Ribo-Seq landscape, with each one detecting on average 85% of them. In contrast, this coverage drops significantly to 38% and 12% for aORFs and iORFs, respectively, strengthening the importance of pooling multiple datasets to characterize pervasive translation events. Relatedly, Figure S7 shows that iORFs are associated with reads in very few samples compared to CDSs which are detected in nearly all samples. However, iORFs are typically short and lowly expressed, rendering their detection in individual samples even more challenging, especially when the sequencing coverage of the experiment is low, as it is the case in numerous datasets analyzed in this study (Figure S6). The lower detection frequency of aORFs across individual samples compared to their overlapping CDSs can be explained by their short size, and provides a mean to partially deconvolute the impact of ORF length on their detection - we recall that they are detected according to their associated-reads irrespective of the frame. Given the inequality of sequencing coverage among samples and the lower detection frequency of aORFs, we do not exclude that the detection of iORFs in individual samples is significantly underestimated. Altogether, these findings show that while using multiple Ribo-Seq datasets is relevant for qualitative analyses aimed at uncovering the diversity of translation signals, caution is warranted for accurate estimation of the detection frequency of lowly expressed short noncoding ORFs. Finally, we examined the robustness of the ORF localization in the global Ribo-Seq landscape with respect to that in the individual ones (Figure 4B). The localization of CDSs and aORFs are highly conserved across the different individual samples, with average Euclidean distances between the global Ribo-Seq landscape and the individual ones much lower than what would be expected by chance (random distribution in grey). This anticipated result reflects the strong optimization of coding region translation to ensure the production of the functional protein. However, it is worth noting that iORF localization is also not random, being significantly conserved in the different experiments compared to the random distribution. Specifically, the localization of intermediate-FIR iORFs, although associated with slightly higher Euclidean distances than the other iORFs, is robust across to the individual samples. Overall, these results suggest that, even pervasive, the translation signals in noncoding regions are consistent in standard conditions.

**Figure 4:**
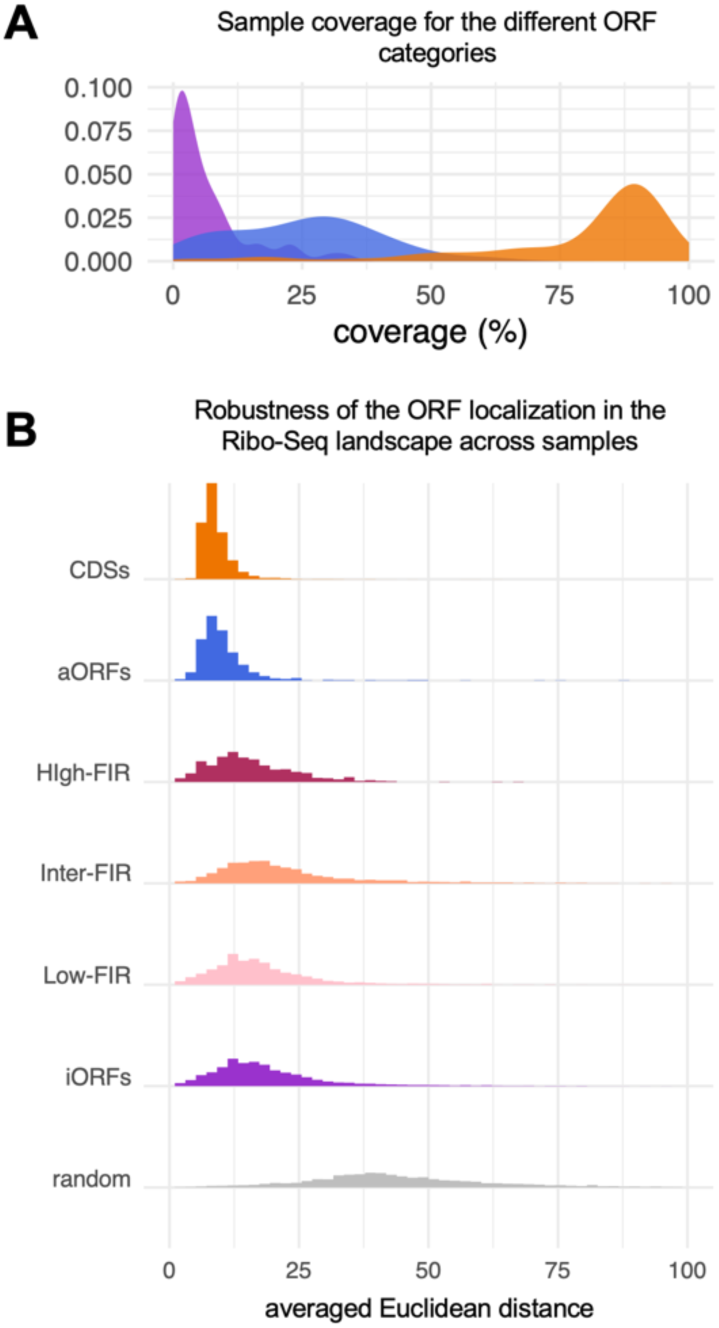
Robustness of translation signals across individual Ribo-Seq samples. **(A)** Distributions of the fractions of ORFs present in the global Ribo-Seq landscape that were found in each sample, with distributions calculated for iORF, aORF, and CDS colored in purple, blue, and orange, respectively. Low coverages indicate that only a small fraction of the ORFs present in the global Ribo-Seq landscape were detected in the corresponding samples, while high coverages reveal that most ORFs of the global Ribo-Seq landscape were found in the individual samples. **(B)** Distributions of the Euclidean distances calculated between the 2D coordinates of an ORF in the global Ribo-Seq landscape and its coordinates in the individual samples (ORFs found in a single sample were excluded from the analysis). Low Euclidean distances reflect ORFs whose coordinates in the individual samples are overall similar, while high distances indicate ORFs that locate in different areas of the individual Ribo-Seq landscapes.

### Rules determining the translation signal in pervasive translation

To elucidate the rules that determine the Ribo-Seq signal of iORFs, we examined their localization within the Ribo-Seq landscape relative to their expression level or several sequence and structural properties. iORFs, irrespective of their corresponding Ribo-Seq landscape area, exhibit comparable expression levels, GC contents, length, foldability potential, and iORFs translated with high specificity are not more conserved across neighboring species than the others (Figure S8). Furthermore, we observed that high-FIR iORFs are slightly more abundant outside UTRs than intermediate-FIR ones, but this effect is small and likely does not account for the overall increase in their translation specificity (one proportion z-test (two-sided), P = 0.003 - Figure S8). Finally, iORFs, regardless of their localization, display comparable amino acid compositions except for Methionine which is significantly enriched in high-FIR iORFs (Figure S9). In fact, it is well acknowledged that the preinitiation complex scans mRNAs from the 5’ searching for ATGs ^62,63^; hosting ATGs is therefore important for being translated with high specificity. This prompted us to characterize the iORF translation initiation codons and systematically depict their impact on the translation specificity (Figure 5A). We detected a start for nearly all high- and intermediate-FIR iORFs (99% and 96%, respectively), while this was only the case for 61% of the low-FIR iORFs, further supporting that although low-FIR ORFs may be associated with translation outcomes, an important fraction of them appear to consist in untranslated ORFs overlapping translated ones. Overall, high-FIR ORFs exhibit a strong tendency to initiate with ATG, or TTG (48% and 10%, respectively, Gini index = 0.75). Conversely, intermediate-FIR iORFs display a wider diversity of start codons, initiating with ATG or six of the nine near-cognate codons with comparable frequencies (Gini index of 0.68) (Figure 5AB) recalling previous reports ^64–66^. In order to investigate how the different near-cognate codons affect translation specificity, we represented the distributions of in-frame read fractions of iORFs relative to their detected start codon (Figure 5C). As expected, initiation at ATG is accompanied by high translation specificity (F0 median = 0.66), while near-cognate codons exhibit a continuum of in-frame read fractions that may reflect different levels of initiation efficiency recalling the observations made in previous studies ^67–72^. In particular, ORFs initiating with NTG codons are associated with higher fractions of in-frame reads than those starting with ANNs (comparison between all merged NTGs versus all ANNs, Mann–Whitney U-test (one-sided), P = 5 × 10^−3^). Finally, to further depict the impact of the genomic context of the start codon on translation specificity, we examined the codon content of the alternative frames of intermediate and high-FIR iORFs. For each intermediate or high-FIR ORF, we calculated the propensity of each codon to be in the ORF frame compared to its two alternative frames (see Methods). Figure 5D shows that high-FIR ORFs are clearly enriched in ATG and TTG codons with respect to their overlapping frames. In contrast, although significantly enriched in ATGs and near-cognate codons compared to their alternative frames, the effect for intermediate-FIR iORFs is less pronounced. These results reveal that hosting ATGs, while important, is not sufficient for being translated with high specificity. Instead, the translation specificity appears to result from the unbalanced distribution of ATGs in the different frames. This supports the hypothesis that, at least in pervasive translation, the translation signals of intermediate specificity reflect translation events occurring in the different RNA frames rather than translation of distinct RNA molecules. We may hypothesize that once associated with the translation machinery, the three RNA frames are in competition for being translated, with the translation of iORFs enriched in ATGs with respect to their alternative frames being more likely to overcome that of their alternative ORFs. Whether additional features (e.g., RNA secondary structure, ribosome binding site, tRNA abundances…) are involved in the translation outcome of iORFs deserves further investigation.

**Figure 5:**
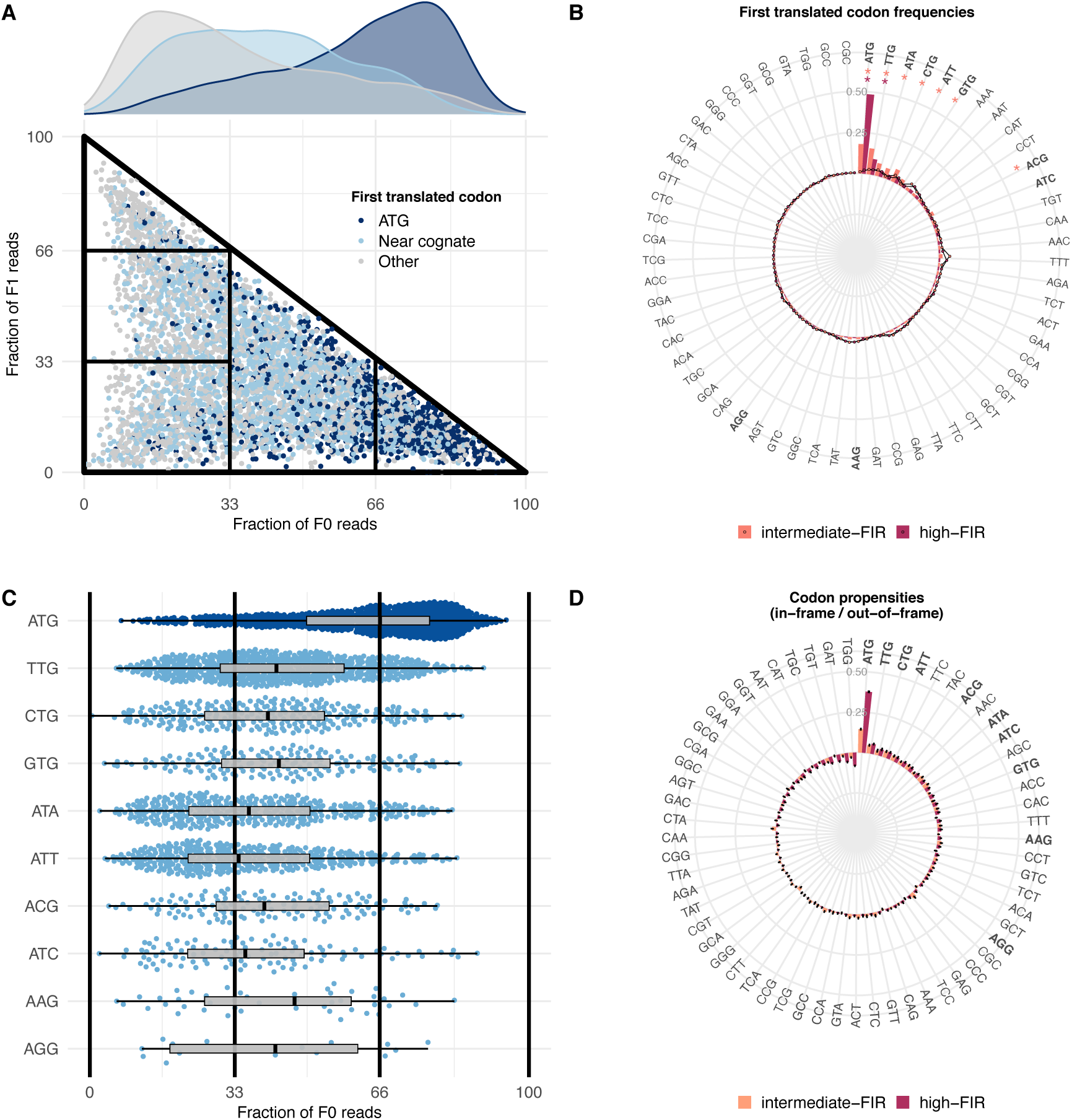
Impact of the initiation codon on the translation specificity in pervasive translation. **(A)** Ribo-Seq landscape of the iORFs for which we detected the translation initiation codon. iORFs are colored according to the type of the first translated codon with dark blue, light blue and grey dots corresponding to initiation with ATG, near cognate codons or other codons, respectively. Densities of the three different types of codons with respect to the fraction of in-frame reads are represented on the top of the plot. **(B)** Barplots representing the frequencies of the 61 codons at the first translated codon for intermediate-FIR (salmon) and high-FIR iORFs (dark red). The small circles (salmon and dark red) indicate the expected frequency of each codon according to its frequency in the intermediate- or high -FIR iORFs, respectively. Codons which are significantly enriched at the first translated position are indicated with a star (one proportion z-test, P < 0.05). Near-cognate codons are represented in bold. **(C)** Distribution of the fraction of in-frame (F0) reads according to the type of the first translated codon. Each dot corresponds to an iORF for which the first translated codon is an ATG (dark blue), or near cognate (light blue). **(D)** Propensities of the 61 codons to be in the frame of the considered iORFs versus in their alternative frames (i.e., out-of-frame). Dark red: propensities of codons to be in the frame of the high-FIR iORFs versus in their alternative frames. Salmon: propensities calculated for intermediate-FIR iORFs. Near cognate codons are represented in bold. Error bars indicate the standard error calculated over 100 random samples of 300 iORFs.

## Discussion

In this work, we characterized the Ribo-Seq landscape of all the ORFs located in ribosome-bound RNAs, thereby revealing the complete palette of translational signals, including canonical and non-canonical ones. We also considered the ORFs with *a priori* no translation outcome in the studied conditions (most of low-FIR ORFs) but which were nevertheless associated with ribosome-protected fragments. Beyond the fact that these ORFs provide a reference to characterize the properties of those that are translated, our data prompt us to hypothesize the existence of a continuum from non-translation outcomes to highly specific translation. In particular, the distinction between untranslated iORFs and iORFs translated with very low specificity can be difficult for technical reasons (coverage and quality of Ribo-Seq data) but also conceptually. Indeed, these translation status may be unstable, and iORFs that are not translated today in standard conditions might be translated in other conditions or in the future. Finally, one should notice that the population of untranslated iORFs is heterogeneous since untranslated iORFs located in ribosome-bound RNAs are more likely to be translated later or in other conditions than those lying in RNAs that do not access the translation machinery, or even worse, those that are not transcribed. In fact, we may hypothesize that if low-FIR iORFs have lost the competition (if any) with their overlapping iORFs for being translated, they nevertheless participate in this competition and, therefore, may play a “passive” role in the translation outcome of the region. All these reasons support the need for characterizing the translation signals of all the ORFs associated with ribosome-protected fragments (i.e., which access to the translation machinery) to delineate the translatome but also the translation potential of a species.

We showed that the translation signals of RNA regions undergoing pervasive translation are much more diverse than previously thought. However, we found that pervasive translation is not random, with pervasive translation signals being qualitatively consistent in standard conditions. Specifically, we showed that iORFs display similar localization in the individual Ribo-Seq landscapes (Figure 4B). We present a topology of the Ribo-Seq landscape of pervasive translation that enables the rational delineation and classification of the entire diversity of Ribo-Seq signals that could be observed in pervasive translation (Figure 6). In particular, we unveiled a twilight zone that is most of the time not considered by classical pipelines which precisely focus on translation signals of high specificity. Our results nevertheless suggest that an important fraction of the ORFs of this region are translated as supported by their corresponding metagenes and MS data. This twilight zone is highly populated in noncoding regions and seems to be a hallmark of pervasive translation. We also highlighted three Ribo-Seq landscape areas of unbalanced ribosome occupancy reflecting RNA regions associated with high translation specificity. These areas are populated in both coding and noncoding regions, however, their status in coding and noncoding regions are not the same. While the translation specificity observed in coding regions mostly results from the regulation of CDS translation to ensure successful protein synthesis, translation specificity in noncoding regions is mainly due to both the codon used to initiate translation and its surrounding genomic context. Furthermore, if the translation status of CDSs is expected to be long-lasting throughout evolution, the one of iORFs, even if specifically translated, is expected to be less stable. Upon mutation, high-FIR ORFs may become poorly translated or untranslated and vice-versa. In other words, the specific translation of noncoding regions is a property that is “innate” and probably short-lived, contrary to the “acquired” and stable translation specificity of CDSs.

**Figure 6:**
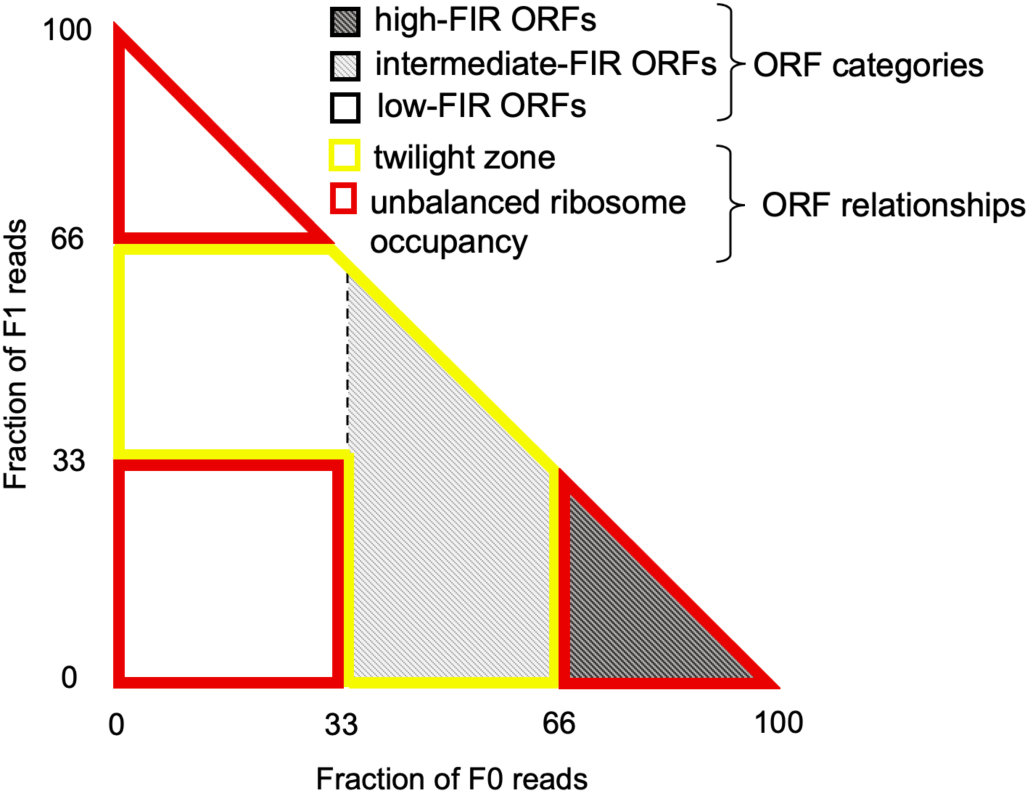
Topology of the yeast Ribo-Seq landscape. Schematic representation of the whole diversity of translation signals that allows (i) their comprehensive and comprehensible classification, and (ii) the efficient and clear identification of areas that are either specific to pervasive translation or present in both pervasive and regular translation.

Indeed, our findings suggest that overall, the high translation specificity of an iORF does not result from the translation optimization of a functional product. Specifically translated iORFs are not characterized by sequence or structural features usually indicative of the emergence of function but instead display higher enrichments in ATGs and TTGs relative to their alternative frames. Whether iORFs enriched in ATGs with respect to their overlapping counterparts are more likely to reach the protein state and, therefore, to be fixed as novel coding products deserve further investigation. Indeed, our MS data showed that a peptide could be detected for aORFs and iORFs associated with different Ribo-Seq landscape areas, strengthening the complex relationship between the specificity of translation and the likelihood of being detected with MS. Moreover, we showed that aORFs for which a peptide was detected with MS were associated with higher numbers of reads regardless of their translation specificity (Figure S2-S3), revealing, this time, the complex relationship between the quantity and the specificity of translation. Finally, leading to a detectable product differs from being functional and selected, though the latter necessarily involves being produced at the protein level. Overall, these results show that the relationship between the expression features of an ORF (quantity and specificity), its capacity to reach the protein state, and its potential to be fixed is still unclear and deserves to be carefully investigated.

The fact that high-FIR iORFs do not distinguish themselves from intermediate or low-FIR iORFs according to features indicative of the emergence of function suggests that most pervasively translated products are not functional (at least in the traditional sense, i.e., not under selection). Nonetheless, a fraction of them may play a biological role in specific or ephemeral conditions like the evolutionarily transient and short-lived sequences reported in Wacholder et al. (2021). It is worth noting that playing a biological role is different from simply having a biochemical activity since the latter does not necessarily involve the former. The distinction between being functional and playing a biological role is even more difficult and not clear conceptually. In fact, the distinction may lie in the timescale, since we usually refer to functional sequences, those that are under selection, i.e., whose function and the conditions under which they are functional persist sufficiently during evolution that we are able to detect evolutionary traces of selection. Here, we propose that iORFs and pervasive translation are functional “collectively” providing the cell with the raw material for selection, of which, from time to time, a handful of functional products (i.e., that will last throughout evolution) may emerge as illustrated with the dozen of candidates identified in Wacholder et al. (2021). This collective function that is mainly related to a process rather than to individual ORFs, precisely relies on the quantity and diversity of the reservoir of freely evolving sequences that can be translated by pervasive translation. In line with Guerra-Almeida and Nunes-da-Fonseca (2020)^5^ these results call for revisiting the concept of function in the context of the emergence of novel coding products. The landscape and diversity of pervasive translation under stress conditions or in tissues associated with emergence of genetic novelty (e.g., testis or brain ^73–76)^, therefore deserves to be carefully investigated. Our representation framework precisely enables large-scale and quantitative analyses of the dynamics of the Ribo-Seq landscape of a species of interest upon condition changes or mutations and, hence, opens the way for future exciting comparative analyses.

## Methods

### Extraction of CDSs, aORFs and noncoding ORFs

The CDSs and noncoding ORFs were extracted with our program ORFtrack ^77^ from the genome of *S. cerevisiae* S288C strain based on the genome annotation of the Saccharomyces Genome Database ^78^. iORFs and aORFs are defined from STOP-to-STOP with a minimal size of 60 nucleotides. We annotated as iORFs, those which do not overlap any annotated feature (i.e., tRNA, rRNA, pseudogene…) of *S. cerevisiae* genome by more than 5% of their length and as aORFs, those whose overlap with a CDS is higher than 5% of the aORF’s length.

### Ribosome profiling analyses

From the Sequence Read Archive ^79^, we manually collected 101 Ribo-Seq runs of wild-type *S. cerevisiae* (strain S288C or BY4741) that were publicly available and realized in standard conditions (see Supplementary Data 1 for the complete list). The independent runs of the technical replicates were pulled together leading to 89 biological datasets. For the treatment and the analysis of the Ribo-Seq raw data, we used our own pipeline ORFribo that we release with the new version of the ORFmine package (see Figure S1 for the detailed pipeline). We first removed the adaptor sequences from the NGS reads using cutadapt ^80^. The trimmed reads were divided into kmers ranging from 25 to 35 nucleotides and were mapped to the CDSs of *S. cerevisiae* using both HISAT2 ^81^ and Bowtie2 ^82^. The position of the ribosome’s P-site was estimated for every kmer using riboWaltz ^83^ and the number of in-frame and out of-frame reads per CDS was computed subsequently. For every dataset, we kept only the kmers for which the distribution of the in-frame reads per CDS was associated with a median value higher than 70% (see Supplementary Data 1 for the retained kmers and Figure S10 for the quality controls), thereby leading to 58 datasets with at least one kmer that fulfills this condition. All the retained kmers, regardless of the dataset they came from, were then pooled together and realigned on the whole genome of *S. cerevisiae.* The total number of in-frame and out-of-frame reads per ORF was subsequently computed for each iORF and aORF.

### Calculations of metagenes

Metagenes were calculated for different subsets of ORFs (e.g., intermediate-, high-FIR ORFs…) as follows: for each ORF of a given subset, we summed over the 58 Ribo-Seq datasets, the number of reads associated with each nucleotide (i.e., reads whose predicted P-site corresponds to the position of the considered nucleotide). We then averaged the count of reads per nucleotide position over all the ORFs of a given subset. The resulting metagene, therefore, represents the read number average per nucleotide. Since codons are constituted by three consecutive nucleotides, ORFs that are translated are expected to be associated with a good triplet periodicity with reads accumulating at every first nucleotide of its in-frame codons. Accumulation of reads at the second or third positions of its in-frame codons rather reflects the translation of the alternative frames +1 or +2, respectively.

### Calculation of sequence and structural properties of the peptides potentially encoded in noncoding ORFs

The HCA foldability score was calculated using the pyHCA tool ^84–87^ integrated in our program ORFold ^77^. We estimated the evolutionary ages of all *S. cerevisiae*’s CDS by phylostratigraphy using ORFdate. Therefore, we performed a BLASTp ^88^ search with the annotated CDS of *S. cerevisiae* (i.e., the focal species) against the complete genomes of 10 saccharomyces (see Figure S11 for the complete list and the associated phylogenetic tree). We only considered BLASTp high scoring pairs (HSPs) with an e-value less than 0.001 and a minimum query coverage of 70%. For each CDS, we identified the more distant species in the phylogenetic tree with respect to the focal species that is associated with a positive HSP match. Horizontal gene transfers are rare events in Eukaryotes; consequently, we consider that the last node shared by the focal and the more distant species with a positive HSP reflects the level of conservation of the considered CDS. This node is therefore used to estimate the CDS’s age according to TimeTree ^89,90^ (Figure S11). As we are interested in the early ages of CDS, the node shared by *S. cerevisiae* and *S. pombe* (the more distant species in the tree with respect to *S. cerevisiae*) is considered as the upper limit for the age estimation, and all CDSs with a match in *S. pombe* are associated with the same upper bounded estimated age (i.e. the one of the last node of the tree) regardless of the fact that the CDS could have additional matches with more distant species outside the tree. We then estimated the age of the iORFs focusing on the Saccharomyces *sensu stricto* (Figure S11) since the latter evolve faster than CDSs. BLASTp were performed with the iORFs of *S. cerevisiae* (focal) as queries and those of the Saccharomyces *sensu stricto* species as targets using the same parameters. The remaining steps for age estimation were the same as those used for CDSs.

The propensity of each codon in noncoding ORFs for being in the frame of the ORF compared to being in its alternative frames is defined as the log ratio of the frequency of the codon in the frame of the ORF versus its frequency in the two alternative frames of the ORF (-+ 30 nucleotides around the considered ORF). It is calculated as follows:

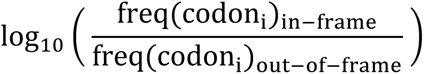

### Detection of the first translated codon

Usually, the translation start is characterized by a noticeable peak of in-frame reads. Therefore, we counted for each codon its number of in-frame reads. Since in noncoding regions, the read coverage is usually low as well as the number of reads per codon, we considered as the translation initiation site, the first 5’ peak of at least 5 reads. Therefore, starting from the 5’ extremity of the ORF, we searched for the first codon *i* associated with at least 5 reads and for which the codons at the positions *i-1* and *i+1* had a number of reads inferior to codon *i*.

### Statistical analyses

All statistical analyses were performed in R (4.0.3) ^91^. To circumvent the P-value problem inherent to large samples ^92^, tests were performed iteratively 1000 times on samples of 300 individuals randomly chosen from the initial sample when it was larger than 300 individuals. The averaged P-value over the 1000 iterations was subsequently calculated.

### Mass Spectrometry Experiments

Yeast strain BY4742 was grown in two different conditions, either in complete synthetic media or in complete synthetic media with MG132 proteasome inhibitor. Cells were lysed in an extraction buffer containing 20mM Hepes, 110mM KoAc, 10 mM MnCl2, 0.5% triton, 0.1% tween and proteases inhibitors. Lysates were cleared by centrifugation and loaded on a gradient 4% to 12% SDS-PAGE. Proteins were extracted from the gels and analyzed by mass spectrometry. For each condition, three gel fractions (1-15 kDa, 15-35 kDa and >35 kDa) were cut in bands of about 2mm and subjected to in-gel trypsin digestion as previously described ^93^ before submission to mass spectrometry analysis. Trypsin631 generated peptides from the three separation regions of the gels were analyzed separately by nanoLC– MSMS using a nanoElute liquid chromatography system (Bruker) coupled to a timsTOF Pro mass spectrometer (Bruker). Peptides were loaded on an Aurora analytical column (ION OPTIK, 25 cm × 75 m, C18, 1.6 m) and separated with a gradient of 0–35% of solvent B for 100 minutes. Solvent A was 0.1% formic acid and 2% acetonitrile in water and solvent B was acetonitrile with 0.1% formic acid. MS and MS/MS spectra were recorded from m/z 100 to 1700 with a mobility scan range from 0.6 to 1.4 V s/cm2. MS/MS spectra were acquired with the PASEF (Parallel Accumulation Serial Fragmentation) ion mobility-based acquisition mode using a number of PASEF MS/MS scans set as 10. MS and MSMS raw data were processed and converted into mgf files with DataAnalysis software (Bruker). Protein identifications were performed using the MASCOT search engine (Matrix Science, London, UK) against coding and noncoding homemade databases 94. Database searches were performed using trypsin cleavage specificity with two possible missed cleavages. Carbamidomethylation of cysteines was set as fixed modification and oxidation of methionines as variable modification. Peptide and fragment tolerances were set at 10 ppm and 0.05 Da, respectively.

### MS Statistical validation of peptides resulting from iORF translation

Peptides identified from noncoding sequences were validated by combining two FDR-controlling procedures: Target-Decoy Competition (TDC) ^95^ and the Benjamini-Hochberg framework (BH) ^96^. Peptide Spectrum Matches (PSM) datasets were first filtered by TDC at 1.7% FDR (coding and statistics). The total non-filtered PSMs were exported and PSM mascot scores were converted into p652 values (p=10-S/10). The p-values were then adjusted with BH procedure and PSM were filtered at 2.5 % FDR (approximately the same number of PSM were validated by TDC and BH at 1.7% and 2.5% FDR, respectively). Finally, in order to ensure a PSM list of candidates with very high confidence (very low FDR), only PSMs validated by both methods were considered.

## Data access

Raw MS data are currently being deposited on PRIDE (https://www.ebi.ac.uk/pride/).

Raw and calculated data used in this study are available as Supplemental Data files except the 101 Ribo-Seq datasets that can be downloaded from the Sequence Read Archive database (Leinonen et al. 2011) (the Accession codes for every dataset are provided in the Table available as Supplementary Data 1). The extraction of all ORFs (CDSs, aORFs and iORFs), the analysis of their sequence and structural properties (foldability potential, ORF conservation) along with their translation activity were calculated using our in-house programs (ORFtrack, ORFold, ORFdate and ORFribo) available in the ORFmine package at: https://github.com/i2bc/ORFmine

All the custom scripts used in this study for the statistical analyses and the generation of Figures 2-5, Figures S2-S10 are available as Supplemental Data files.

## Acknowledgments

Works by CP and PR were supported by French government fellowships. HA and SB works were supported by ANR Actimeth (19-CE12-0004-02) and ANR Rescue Ribosome (17-CE12-0024). This work has benefited from the facilities and expertise of the I2BC proteomic platform (Proteomic-Gif, SICaPS) supported by IBiSA, Ile de France Region, Plan Cancer, CNRS and Paris-Saclay University. Conceptualization: CP, ON, AL; Methodology: CP, HA, NC, SB, DC, ON, AL; Investigation and development: CP, HA, NC, SB, DC, PR, CR, SA, AL; Writing and editing: CP, NC, SB, PR, OL, ON, AL; Supervision: AL.

## Supplemental Material

**Supplemental Table S1.** Count of the number of ORFs (e.g. CDSs, aORFs and iORFs) with more than 50 Ribo-Seq reads in each Ribo-Seq landscape region and a read coverage of at least 30%,.

|  | Low-FIR | Intermediate-FIR | High-FIR | Total |
| --- | --- | --- | --- | --- |
| **CDSs** | 84 | 81 | 5403 | 5568 |
| **aORFs** | 77627 | 680 | 67 | 78374 |
| **iORFs** | 3526 | 2226 | 1036 | 6788 |

**Supplemental Table S2.** Count of the number of ORFs (i.e. CDSs, aORFs and iORFs) with more than 50 Ribo-Seq reads and a read coverage of at least 30%, for which at least one (aORFs and iORFs) or two peptides (CDSs) were observed with MS in the different conditions.

|  | Standard | Inhibited Proteasome | Intersection | Union |
| --- | --- | --- | --- | --- |
| **CDSs** | 3087 | 3639 | 3000 | 3726 |
| **aORFs** | 127 | 152 | 62 | 217 |
| **iORFs** | 15 | 23 | 4 | 34 |

**Supplemental Figure S1:**
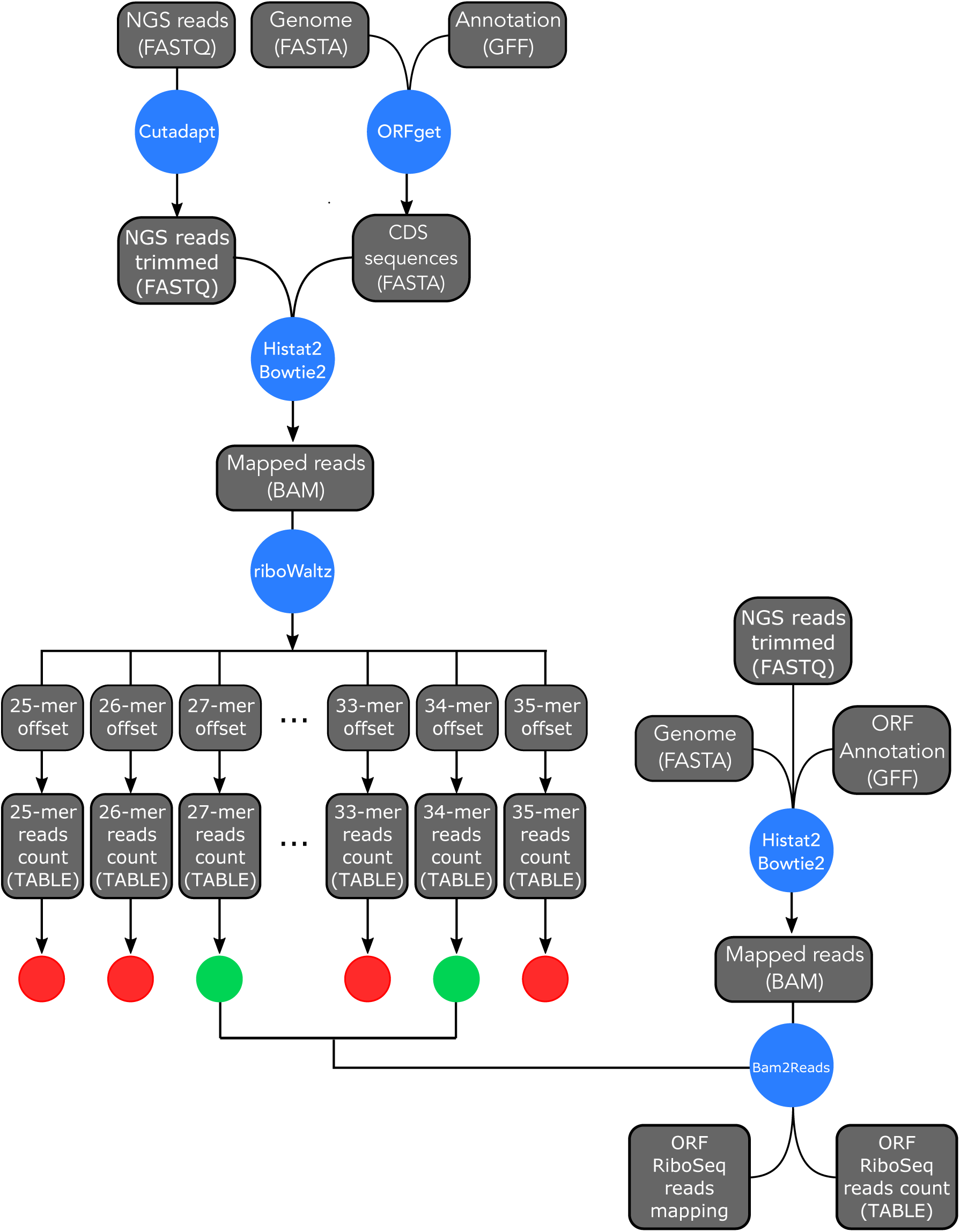
Pipeline of ORFribo, from the novel version of the ORFmine package. For the treatment and the analysis of the Ribo-Seq raw data, we used our own pipeline ORFribo released with the present study. **(Left)** The adaptor sequences from the NGS reads were removed using cutadapt (Martin 2011). Based on the *S. cerevisiae* genome annotation (Cherry et al. 2012) and sequence, we obtained all the intronless nucleotide sequences of the CDSs with our tool ORFget. The trimmed reads were mapped to the CDSs of *S. cerevisiae* using both HISAT2 (Kim et al. 2019) and Bowtie2 (Langmead and Salzberg 2012). The mapped reads were divided into kmers ranging from 25 to 35 nucleotides. The position of the ribosome’s P-site on the reads (called offset) was estimated for every kmer using riboWaltz (Lauria et al. 2018) and the number of in-frame and out-of-frame reads per CDS was computed subsequently (generating reads’ count tables). For every dataset, we kept only the kmers for which the distribution of the in-frame reads per CDS was associated with a median value higher than 70% (represented with green circles) and discarded all the kmers that do not fulfill this requirement (represented with red circles). **(Right)** Then the trimmed reads were mapped this time to the whole *S. cerevisiae* genome including noncoding regions. The number of in-frame and out-of-frame reads per ORF (i.e. iORFs and aORFs) as well as their position along the ORF’s sequence were computed only for the kmers that were retained in the previous step (represented with green circles) by our tool Bam2Reads and using the ORFs’ annotation resulting from our tool ORFtrack (Papadopoulos et al. 2022). For each ORF, the number of reads in its different frames (F0, F1 and F2) is then reported in a table of counts.

**Supplemental Figure S2:**
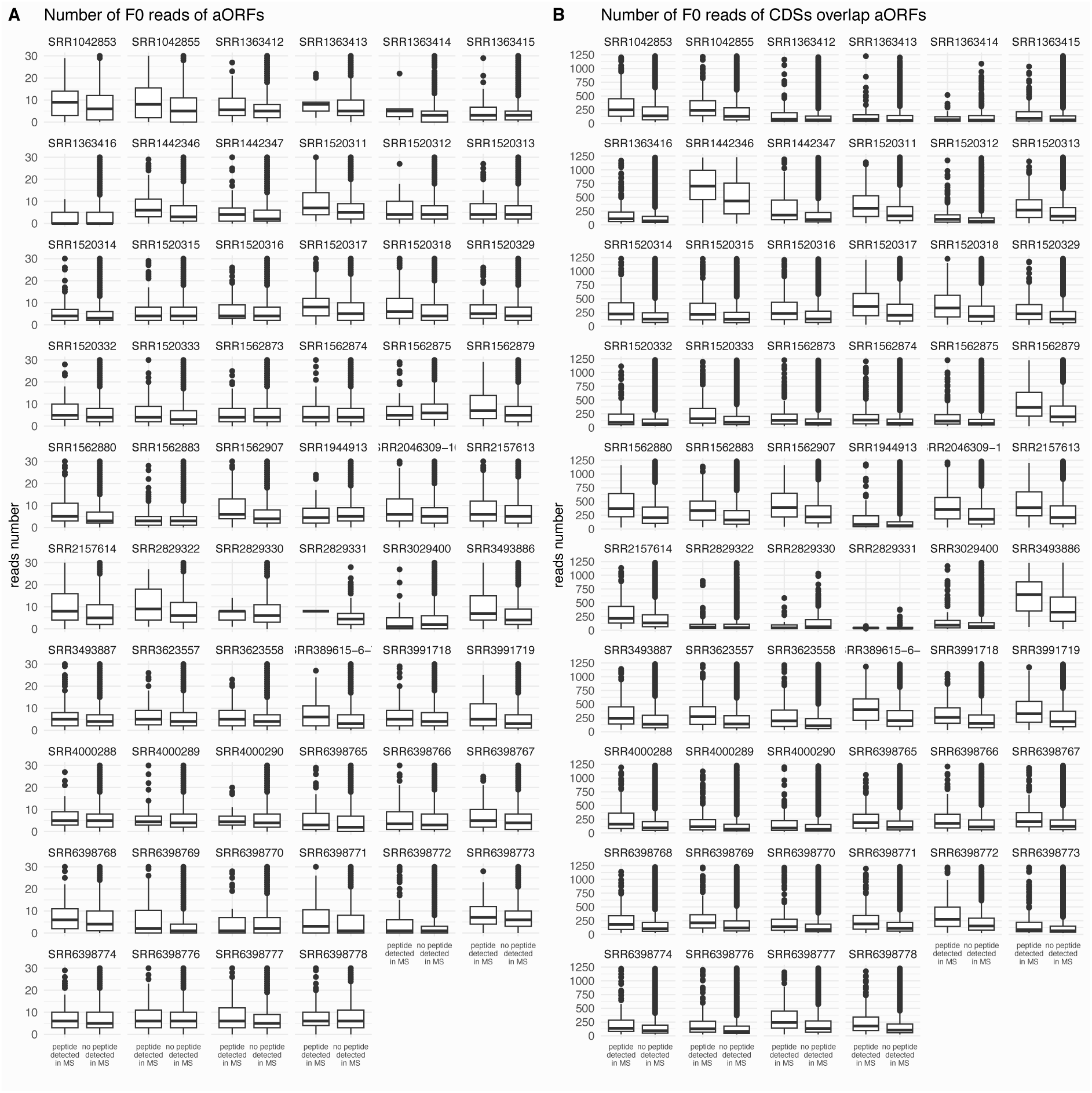
Number of F0 reads of aORFs in individual Ribo-Seq samples. **(A)** Each individual plot corresponds to an individual Ribo-Seq sample. Sample IDs are indicated on the top of each plot. For each plot: number of F0 reads of aORFs for which a peptide was detected in MS experiments (left) and aORFs without any detected peptide (right). **(B)** same as (A) but number of F0 reads of CDSs that overlap aORFs with a MS detected peptide (left) and CDSs whose overlapping aORFs were not associated with a detected peptide (right).

**Supplemental Figure S3:**
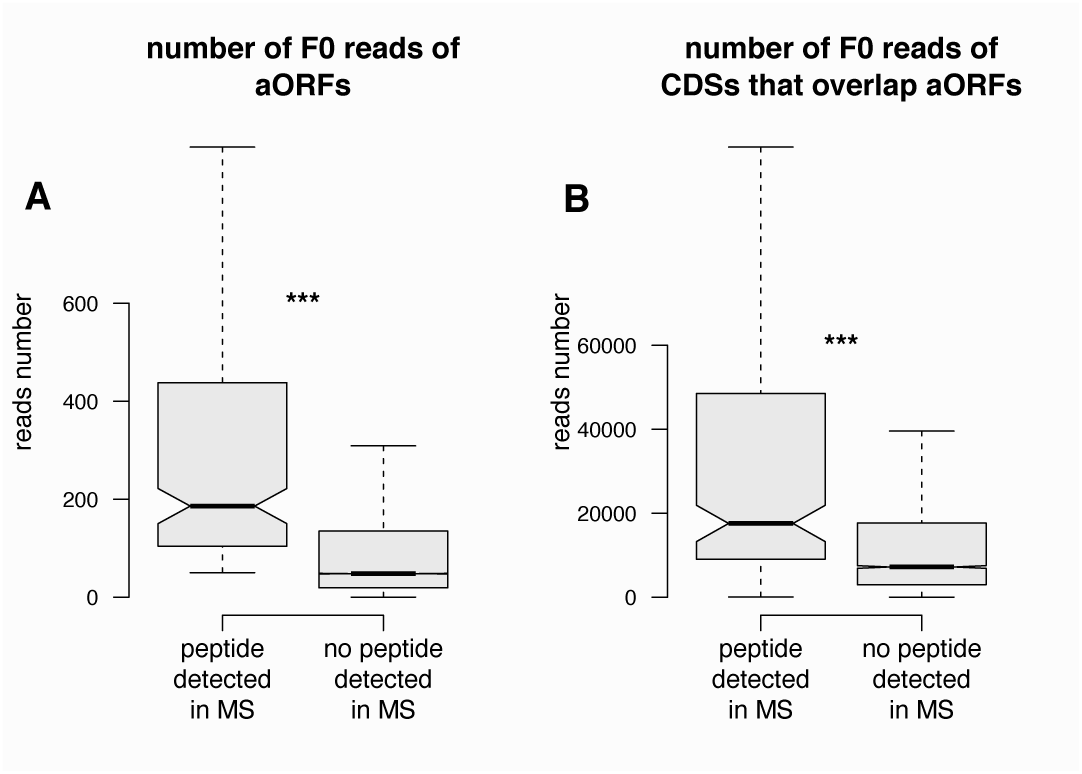
aORFs with a detected MS peptide are associated with higher number of F0 reads. **(A)** number of F0 reads computed over all samples of aORFs for which a peptide was detected in MS experiments (left) and aORFs without any detected peptide (right). **(B)** number of F0 reads over computed all samles of CDSs that overlap aORFs with a MS detected peptide (left) and CDSs whose overlapping aORFs were not associated with a detected peptide (right).

**Supplemental Figure S4:**
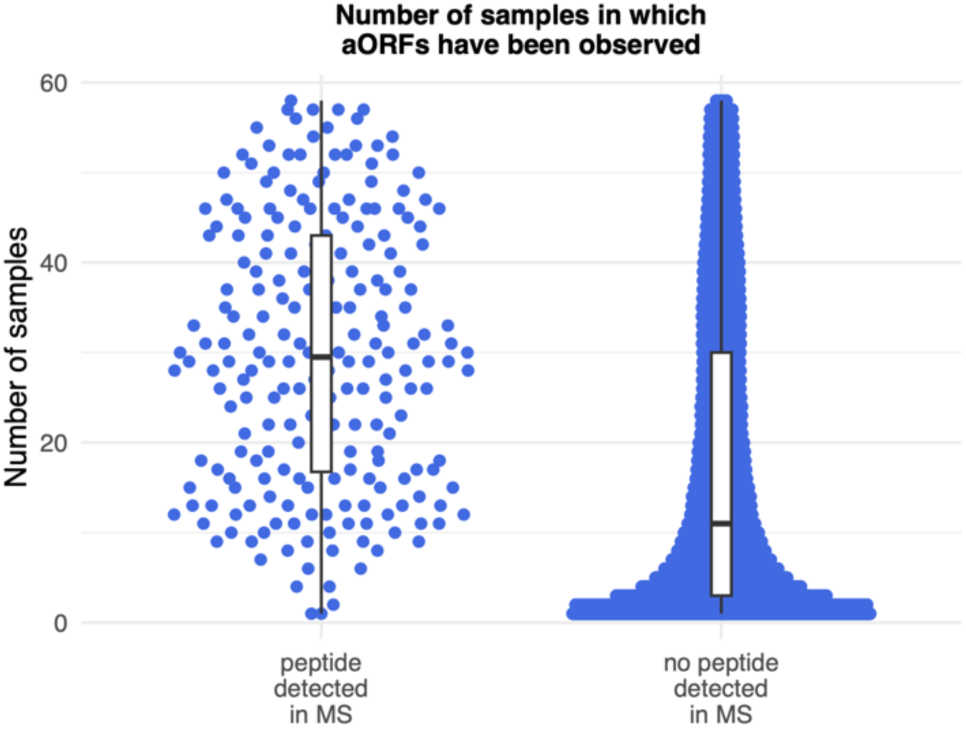
aORFs with a detected MS peptide are detected in more Ribo-Seq samples than the others. Number of Ribo-Seq samples in which an aORFs for which a peptide was detected in MS experiments (left) and aORFs without any detected peptide (right).

**Supplemental Figure S5:**
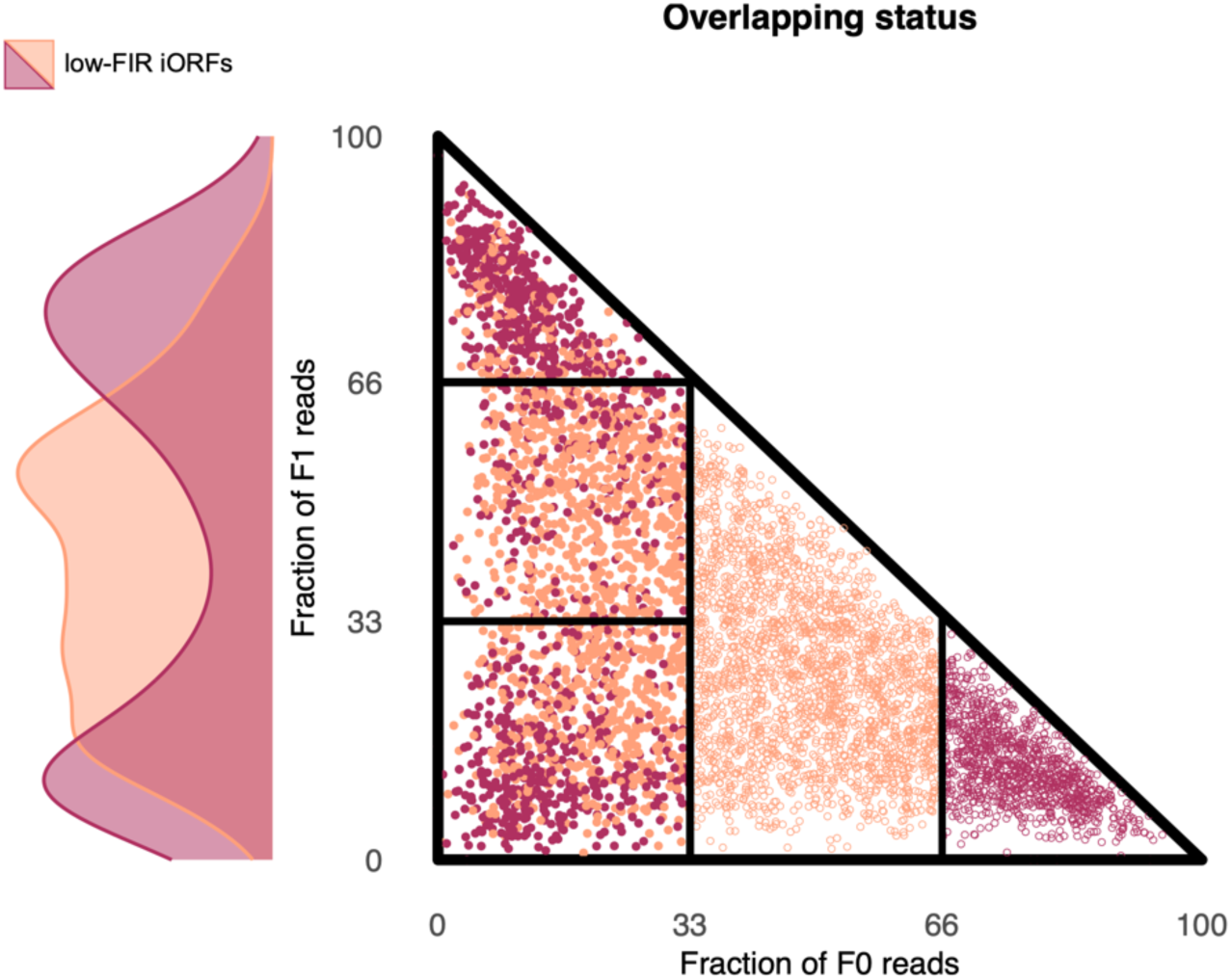
Overlap relationships between read-associated iORFs. Overlap relationships involving high-FIR (empty dark red circles) and intermediate-FIR (empty salmon circles) iORFs with low-FIR iORFs (filled circles). low-FIR iORFs are colored according to the iORF they overlap (dark red or salmon when they overlap with at least one high-FIR or one intermediate-FIR iORF, respectively). The densities of low-FIR ORFs that overlap high-FIR or intermediate-FIR iORF are presented in dark red or salmon respectively on the left part of the graph.

**Supplemental Figure S6_1:**
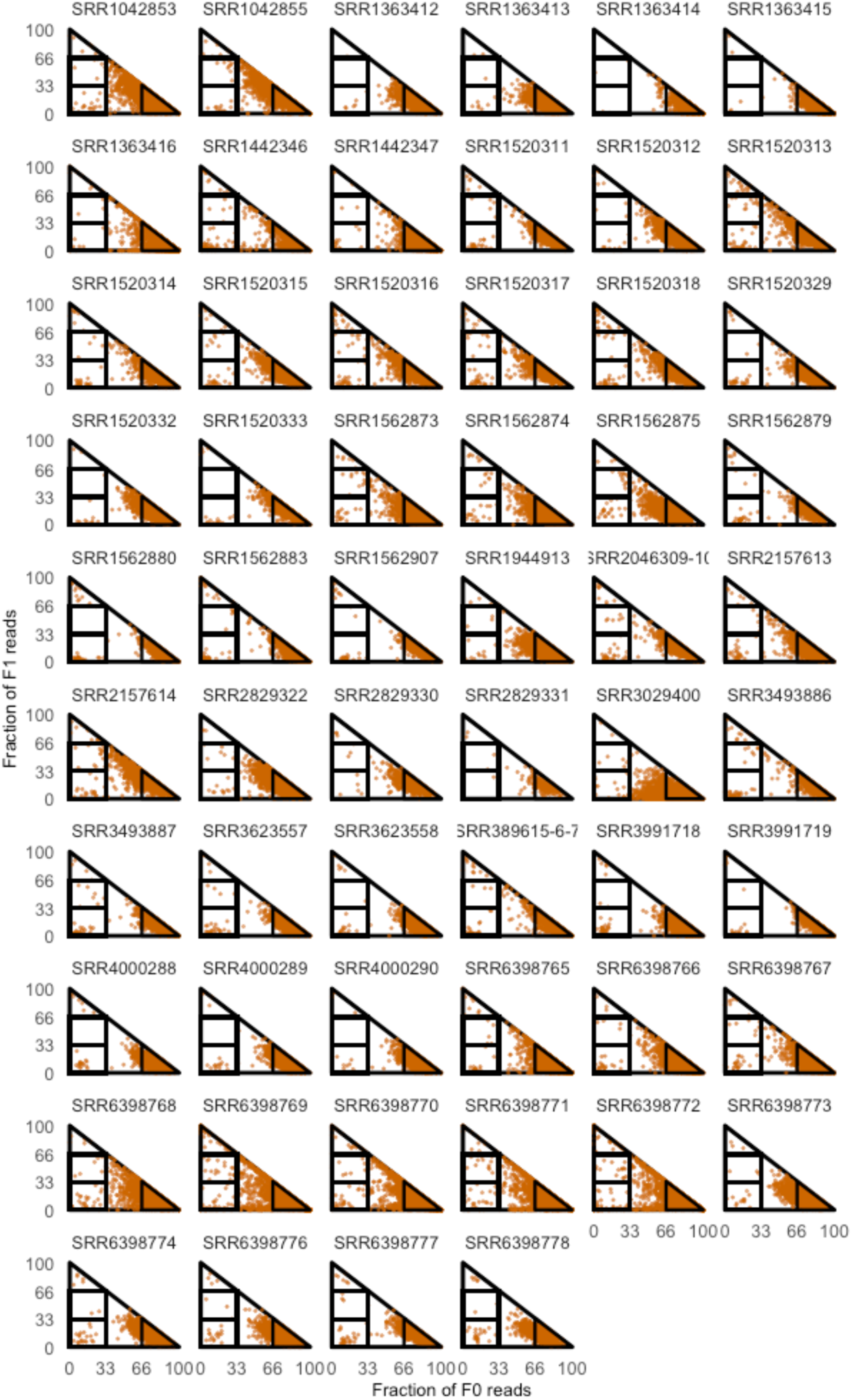
CDS individual Ribo-Seq landscapes calculated for each sample. Sample IDs are indicated on the top of each Ribo-Seq landscape. CDSs are plotted on the Ribo-Seq landscape of a given sample if they are associated with at least 10 reads regardless of the frame.

**Supplemental Figure S6_2:**
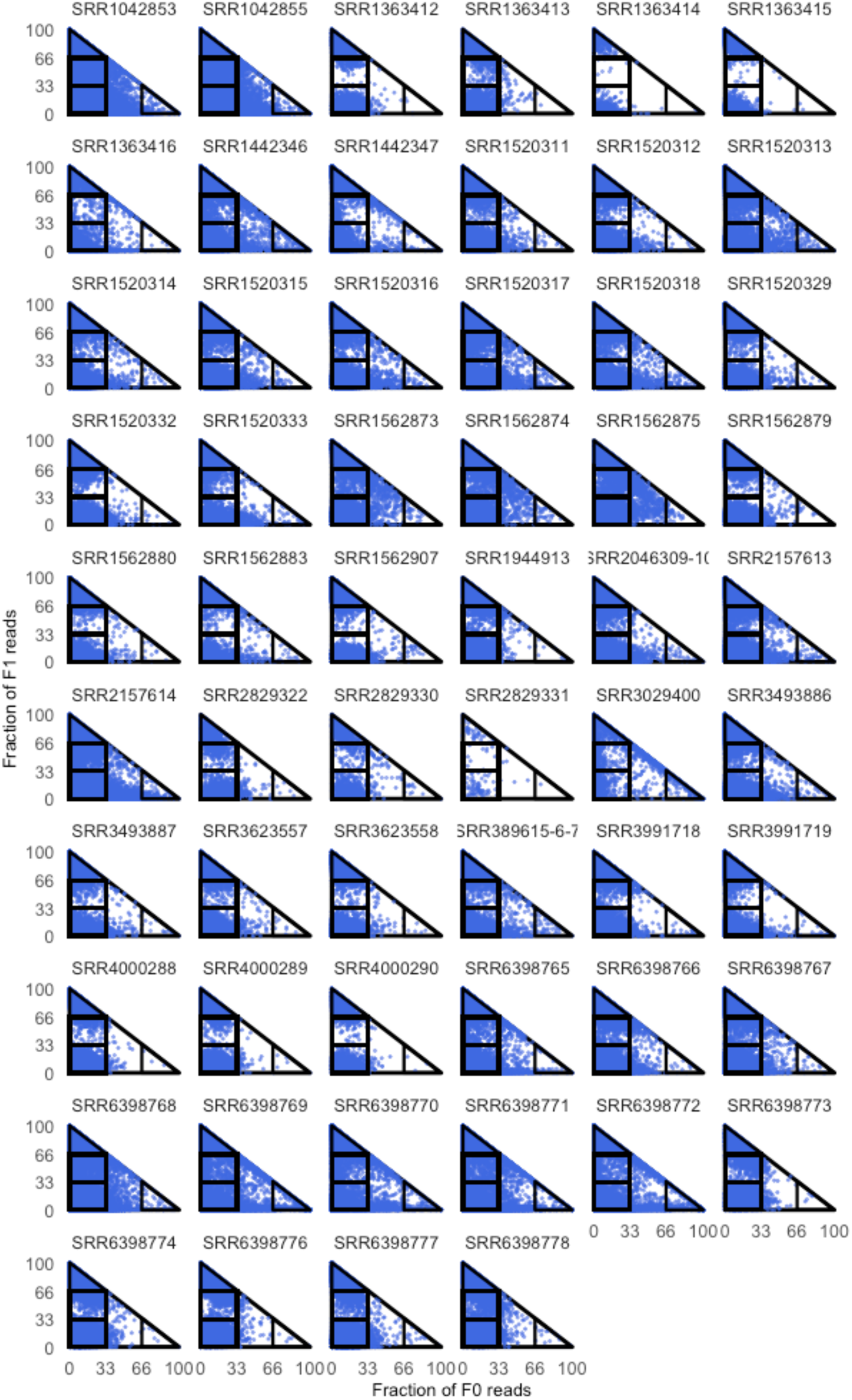
aORF individual Ribo-Seq landscapes calculated for each sample. Sample IDs are indicated on the top of each Ribo-Seq landscape. aORFs are plotted on the Ribo-Seq landscape of a given sample if they are associated with at least 10 reads regardless of the frame.

**Supplemental Figure S6_3:**
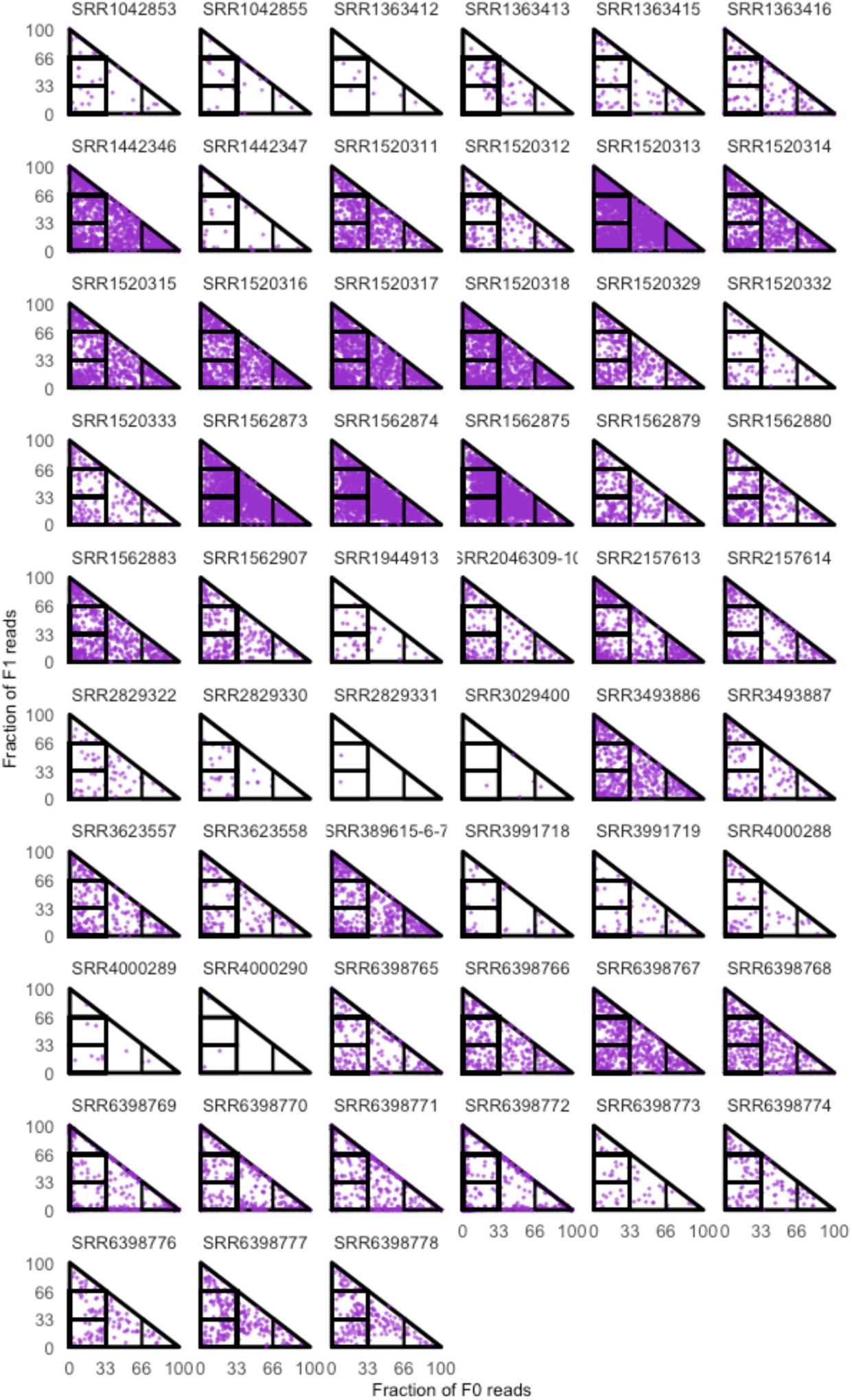
iORF individual Ribo-Seq landscapes calculated for each sample. Sample IDs are indicated on the top of each Ribo-Seq landscape. iORFs are plotted on the Ribo-Seq landscape of a given sample if they are associated with at least 10 reads regardless of the frame.

**Supplemental Figure S7:**
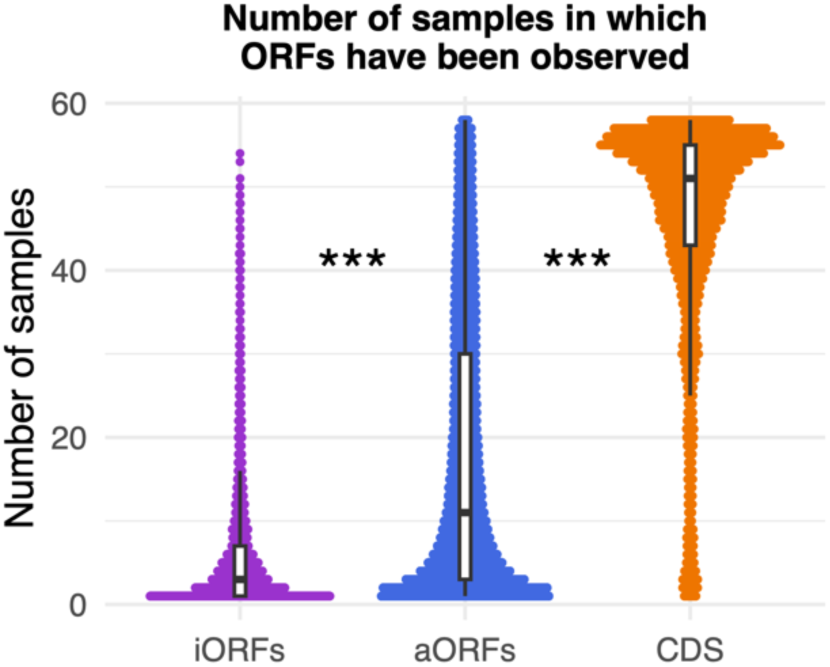
Number of samples in which ORFs have been detected. Distributions of the number of samples in which iORFs, aORFs and CDSs were detected in purple, blue and orange, respectively. ORFs are retained if they are associated with at least 10 reads regardless of the frame. The p-values were computed with a two-sided Kolmogorov–Smirnov test. Asterisks denote level of significance: *p < 5 × 10-2, **p < 1 × 10-2, ***p < 1 × 10-3.

**Supplemental Figure S8:**
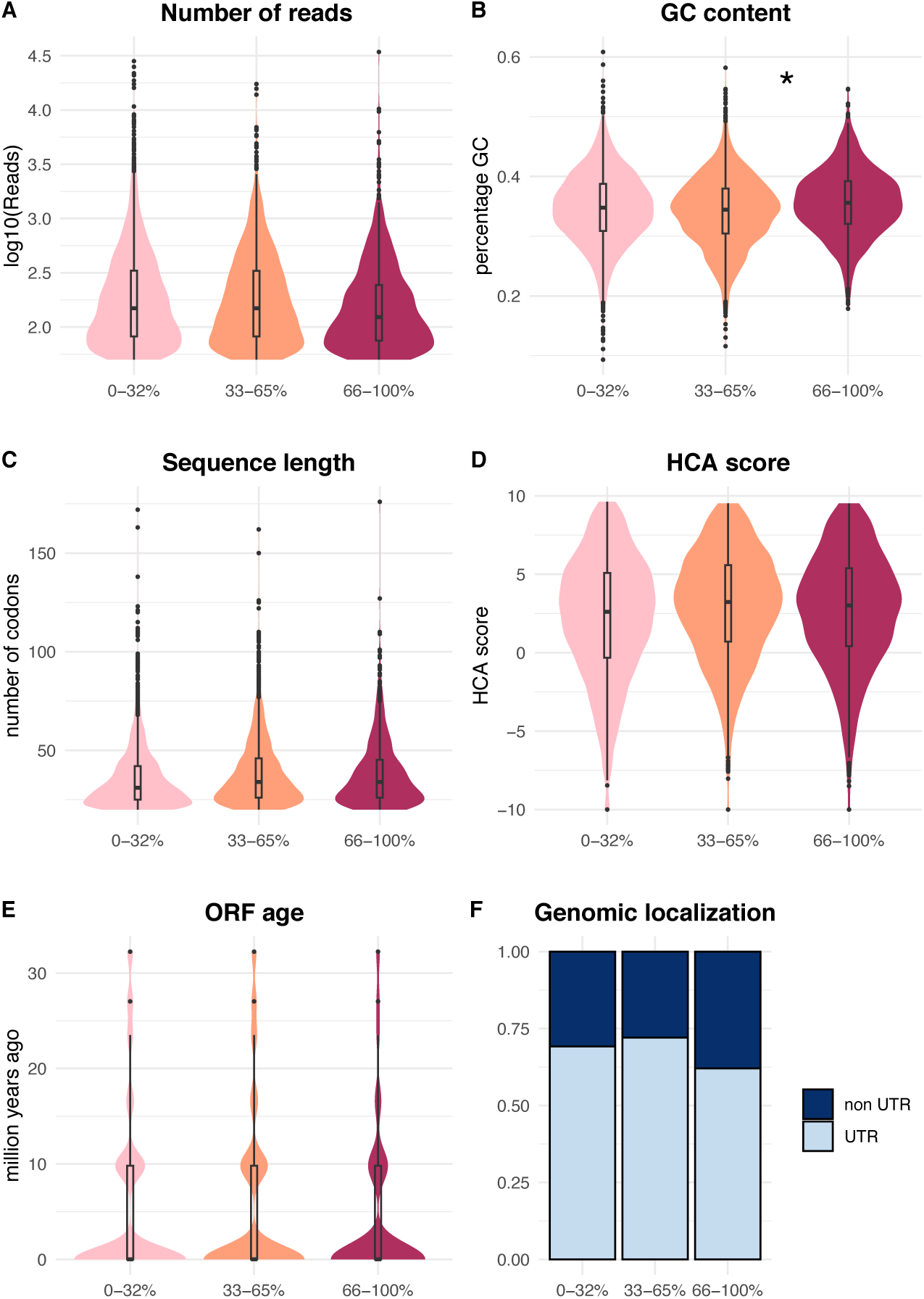
Properties of iORFs according to their translation specificity. Distributions for low-, intermediate-, and high-FIR iORFs of **(A)** the number of reads regardless of the frame, **(B)** GC-content, **(C)** sequence length in amino acids, **(D)** HCA scores calculated on their translated amino-acid sequence, **(E)** estimated evolutionary ages. Low-, intermediate- and high-FIR iORF distributions are represented in light pink, salmon and dark red, respectively. **(F)** Stacked barplots of the genomic localization of low-, intermediate-, and high-FIR iORFs (light and dark blue for iORFs located in UTRs and outside UTRs, respectively). The p-values were computed with a two-sided Kolmogorov–Smirnov test. Asterisks denote level of significance: *p < 5 × 10-2, **p < 1 × 10-2, ***p < 1 × 10-3.

**Supplemental Figure S9:**
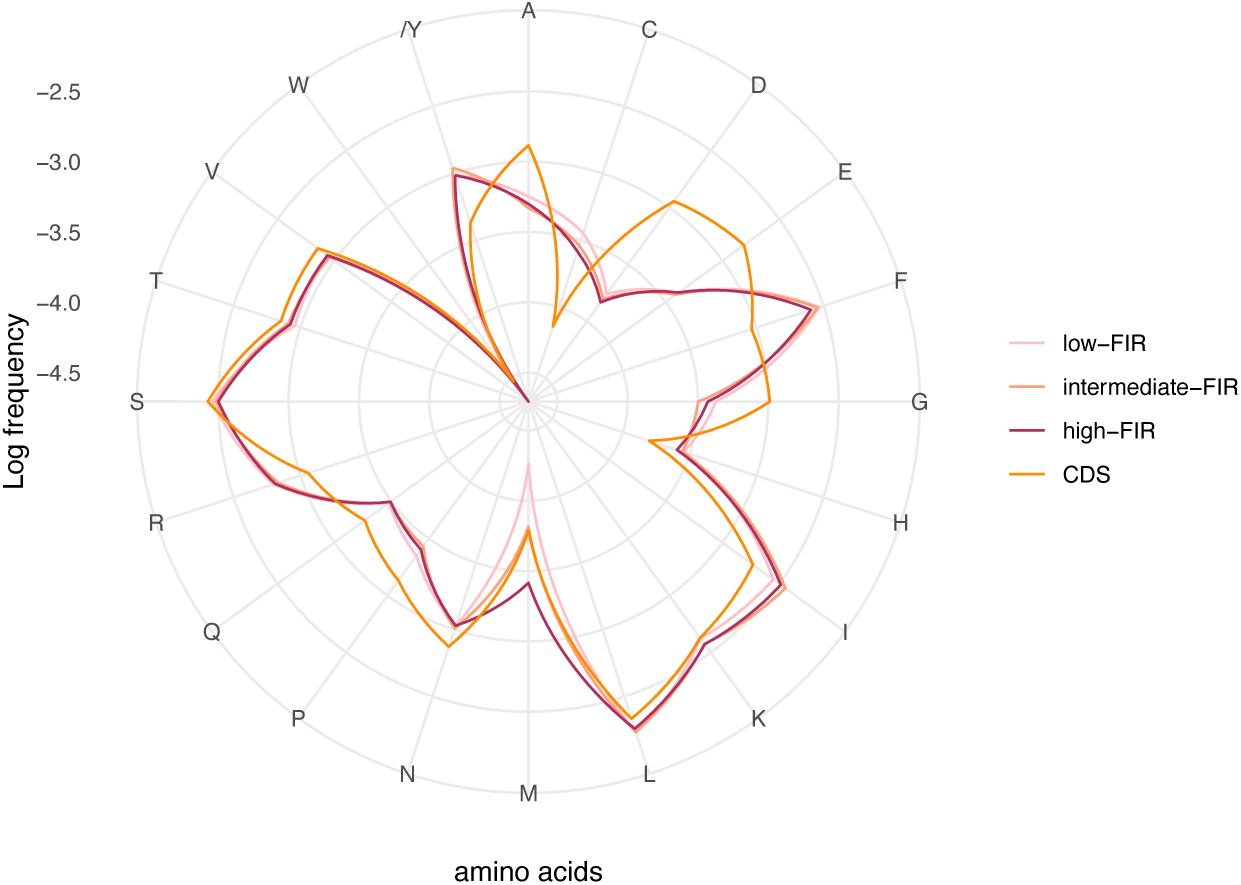
Amino acid compositions of the different iORF categories are distinct from that of CDSs and similar between them, except for Methionine. Radar plot reflecting the 20 amino acid frequencies of low-, intermediate-, high-FIR iORFs and CDSs in light pink, salmon, dark red and orange, respectively.

**Supplemental Figure S10:**
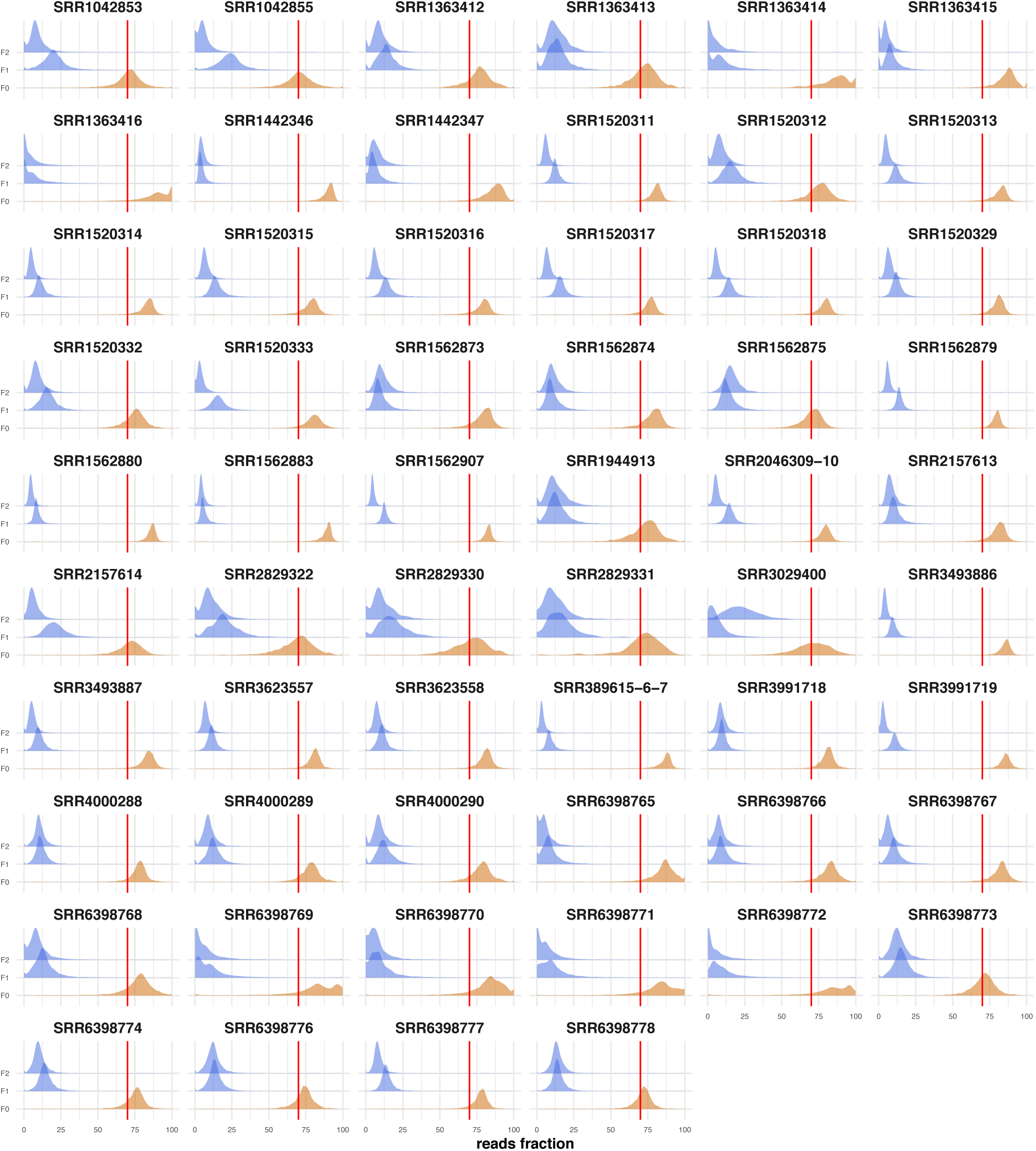
Quality control of the retained Ribo-Seq datasets. Distributions of the fractions of reads in the F0 (orange), F1 (blue), F2 (blue) frames of the annotated CDSs for each dataset. Sample IDs are presented on the top of each plot. The vertical red lines indicate the quality threshold of a median of 70% of reads mapping to the F0 frames of CDSs used as quality control. All CDSs annotated in the GFF file of the well-annotated yeast are used as reference. Although annotation errors may occur, we believe they are rare in *S. cerevisiae* and will have little impact on the quality control step. Specifically, misannotated CDSs are expected to be associated with increases in shifted reads relative to their F0 frame and in the worst scenario, will decrease the overall fraction of F0 reads of the sample (though unlikely), making us wrongly eliminate the corresponding sample.

**Supplemental Figure S11:**
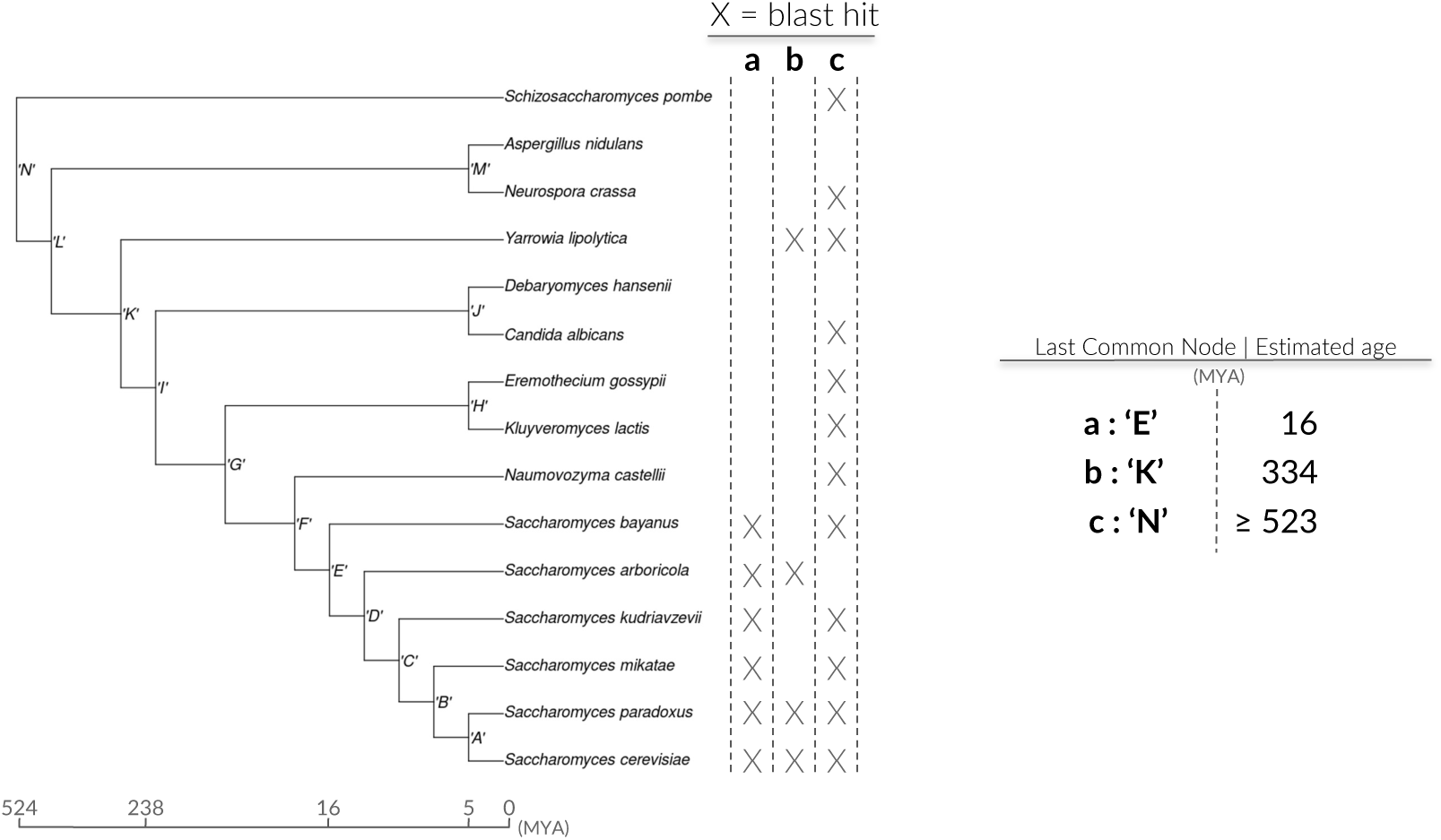
Estimation of ORFs’ ages. Left: phylogenetic tree used to estimate the ages of yeast CDSs. The ages of iORFs were estimated based on the subtree (*S.cerevisiae*, *S. bayanus* - Node E). Each ORF (a,b,c) is screened with BLASTp (Altschul et al. 1990) against each genome of the tree and “X” stands for cases where a hit has been found in the corresponding genomes. The more distant hit with respect to the focal species (here *S. cerevisiae*) is used to assign the last common ancestor (node). Right: the corresponding node is used to estimate the age of the ORF with TimeTree (Hedges et al. 2006; Kumar et al. 2022).

## References

1. Hanada, K., Zhang, X., Borevitz, J. O., Li, W.-H. & Shiu, S.-H. A large number of novel coding small open reading frames in the intergenic regions of the Arabidopsis thaliana genome are transcribed and/or under purifying selection. Genome research 17, 632–640 (2007).

2. Yang, X. et al. Discovery and annotation of small proteins using genomics, proteomics, and computational approaches. Genome research 21, 634–641 (2011).

3. Couso, J.-P. & Patraquim, P. Classification and function of small open reading frames. Nature reviews Molecular cell biology 18, 575–589 (2017).

4. Orr, M. W., Mao, Y., Storz, G. & Qian, S.-B. Alternative ORFs and small ORFs: shedding light on the dark proteome. Nucleic Acids Research 48, 1029–1042 (2020).

5. Guerra-Almeida, D. & Nunes-da-Fonseca, R. Small open reading frames: how important are they for molecular evolution? Frontiers in Genetics 11, 574737 (2020).

6. Guerra-Almeida, D., Tschoeke, D. A. & Nunes-da-Fonseca, R. Understanding small ORF diversity through a comprehensive transcription feature classification. DNA Research 28, dsab007 (2021).

7. Kapranov, P. et al. Large-scale transcriptional activity in chromosomes 21 and 22. Science 296, 916–919 (2002).

8. Ingolia, N. T., Ghaemmaghami, S., Newman, J. R. & Weissman, J. S. Genome-wide analysis in vivo of translation with nucleotide resolution using ribosome profiling. science 324, 218–223 (2009).

9. Clark, M. B. et al. The reality of pervasive transcription. PLoS Biol 9, e1000625 (2011).

10. Carvunis, A.-R., et al. Proto-genes and de novo gene birth. Nature 487, 370–374 (2012).

11. Jensen, T. H., Jacquier, A. & Libri, D. Dealing with pervasive transcription. Molecular cell 52, 473–484 (2013).

12. Chew, G.-L. et al. Ribosome profiling reveals resemblance between long non-coding RNAs and 5′ leaders of coding RNAs. Development 140, 2828–2834 (2013).

13. Aspden, J. L. et al. Extensive translation of small open reading frames revealed by Poly-Ribo-Seq. Elife 3, e03528 (2014).

14. Bazzini, A. A. et al. Identification of small ORFs in vertebrates using ribosome footprinting and evolutionary conservation. The EMBO journal 33, 981–993 (2014).

15. Smith, J. E. et al. Translation of small open reading frames within unannotated RNA transcripts in Saccharomyces cerevisiae. Cell reports 7, 1858–1866 (2014).

16. Ingolia, N. T. et al. Ribosome profiling reveals pervasive translation outside of annotated protein-coding genes. Cell reports 8, 1365–1379 (2014).

17. Ruiz-Orera, J., Verdaguer-Grau, P., Villanueva-Cañas, J., Messeguer, X. & Albà, M. M. Translation of neutrally evolving peptides provides a basis for de novo gene evolution. Nat. Ecol. Evol. 2: 890–896. Nature ecology & evolution (2018).

18. Ruiz-Orera, J. & Albà, M. M. Translation of small open reading frames: roles in regulation and evolutionary innovation. Trends in Genetics 35, 186–198 (2019).

19. Chen, J. et al. Pervasive functional translation of noncanonical human open reading frames. Science 367, 1140–1146 (2020).

20. Blevins, W. R. et al. Uncovering de novo gene birth in yeast using deep transcriptomics. Nature communications 12, 1–13 (2021).

21. Papadopoulos, C. et al. Intergenic ORFs as elementary structural modules of de novo gene birth and protein evolution. Genome Research 31, 2303–2315 (2021).

22. 22. Wacholder, A., Acar, O. & Carvunis, A.-R. A reference translatome map reveals two modes of protein evolution. BioRxiv (2021).

23. Smith, C. et al. Pervasive translation in Mycobacterium tuberculosis. eLife 11, e73980 (2022).

24. Parikh, S. B., Houghton, C., Van Oss, S. B., Wacholder, A. & Carvunis, A. Origins, evolution, and physiological implications of de novo genes in yeast. Yeast 39, 471–481 (2022).

25. Mudge, J. M. et al. Standardized annotation of translated open reading frames. Nature Biotechnology 40, 994–999 (2022).

26. Slavoff, S. A. et al. Peptidomic discovery of short open reading frame–encoded peptides in human cells. Nature chemical biology 9, 59 (2013).

27. Prabakaran, S. et al. Quantitative profiling of peptides from RNAs classified as noncoding. Nature communications 5, 1–10 (2014).

28. Ma, J. et al. Discovery of human sORF-encoded polypeptides (SEPs) in cell lines and tissue. Journal of proteome research 13, 1757–1765 (2014).

29. Hsu, P. Y. & Benfey, P. N. Small but mighty: functional peptides encoded by small ORFs in plants. Proteomics 18, 1700038 (2018).

30. Cao, X. et al. Comparative Proteomic Profiling of Unannotated Microproteins and Alternative Proteins in Human Cell Lines. J. Proteome Res. 19, 3418–3426 (2020).

31. Cuevas, M. V. R. et al. Most non-canonical proteins uniquely populate the proteome or immunopeptidome. Cell reports 34, 108815 (2021).

32. Zheng, E. B. & Zhao, L. Protein evidence of unannotated ORFs in Drosophila reveals diversity in the evolution and properties of young proteins. eLife 11, e78772 (2022).

33. Begun, D. J., Lindfors, H. A., Kern, A. D. & Jones, C. D. Evidence for de Novo Evolution of Testis-Expressed Genes in the Drosophila yakuba/Drosophila erecta Clade. Genetics 176, 1131–1137 (2007).

34. Levine, M. T., Jones, C. D., Kern, A. D., Lindfors, H. A. & Begun, D. J. Novel genes derived from noncoding DNA in Drosophila melanogaster are frequently X-linked and exhibit testis-biased expression. Proceedings of the National Academy of Sciences 103, 9935–9939 (2006).

35. Cai, J., Zhao, R., Jiang, H. & Wang, W. De novo origination of a new protein-coding gene in Saccharomyces cerevisiae. Genetics 179, 487–496 (2008).

36. Zhou, Q. et al. On the origin of new genes in Drosophila. Genome research 18, 1446– 1455 (2008).

37. Knowles, D. G. & McLysaght, A. Recent de novo origin of human protein-coding genes. Genome research 19, 1752–1759 (2009).

38. Siepel, A. Darwinian alchemy: Human genes from noncoding DNA. Genome research 19, 1693–1695 (2009).

39. Tautz, D. & Domazet-Lošo, T. The evolutionary origin of orphan genes. Nature Reviews Genetics 12, 692–702 (2011).

40. Wu, D.-D., Irwin, D. M. & Zhang, Y.-P. De novo origin of human protein-coding genes. PLoS genetics 7, e1002379 (2011).

41. Wissler, L., Godmann, L. & Bornberg-Bauer, E. Evolutionary dynamics of simple sequence repeats across long evolutionary time scale in genus Drosophila. Trends in Evolutionary Biology 4, e7–e7 (2012).

42. Murphy, D. N. & McLysaght, A. De novo origin of protein-coding genes in murine rodents. PloS one 7, e48650 (2012).

43. Zhao, L., Saelao, P., Jones, C. D. & Begun, D. J. Origin and spread of de novo genes in Drosophila melanogaster populations. Science 343, 769–772 (2014).

44. Schlötterer, C. Genes from scratch–the evolutionary fate of de novo genes. Trends in Genetics 31, 215–219 (2015).

45. Bornberg-Bauer, E., Schmitz, J. & Heberlein, M. Emergence of de novo proteins from ‘dark genomic matter’by ‘grow slow and moult’. Biochemical Society Transactions 43, 867–873 (2015).

46. Li, Z.-W. et al. On the origin of de novo genes in Arabidopsis thaliana populations. Genome biology and evolution 8, 2190–2202 (2016).

47. Wilson, B. A., Foy, S. G., Neme, R. & Masel, J. Young genes are highly disordered as predicted by the preadaptation hypothesis of de novo gene birth. Nature ecology & evolution 1, 1–6 (2017).

48. Gubala, A. M. et al. The goddard and saturn genes are essential for Drosophila male fertility and may have arisen de novo. Molecular biology and evolution 34, 1066–1082 (2017).

49. Vakirlis, N. et al. A molecular portrait of de novo genes in yeasts. Molecular biology and evolution 35, 631–645 (2018).

50. Schmitz, J. F., Ullrich, K. K. & Bornberg-Bauer, E. Incipient de novo genes can evolve from frozen accidents that escaped rapid transcript turnover. Nature ecology & evolution 2, 1626–1632 (2018).

51. Van Oss, S. B. & Carvunis, A.-R. De novo gene birth. PLoS genetics 15, (2019).

52. Zhang, L. et al. Rapid evolution of protein diversity by de novo origination in Oryza. Nature ecology & evolution 3, 679–690 (2019).

53. Prabh, N. & Rödelsperger, C. De novo, divergence, and mixed origin contribute to the emergence of orphan genes in pristionchus nematodes. G3: Genes, Genomes, Genetics 9, 2277–2286 (2019).

54. Vakirlis, N. et al. De novo emergence of adaptive membrane proteins from thymine-rich genomic sequences. Nature communications 11, 1–18 (2020).

55. Vakirlis, N., Carvunis, A.-R. & McLysaght, A. Synteny-based analyses indicate that sequence divergence is not the main source of orphan genes. eLife 9, (2020).

56. Heames, B., Schmitz, J. & Bornberg-Bauer, E. A continuum of evolving de novo genes drives protein-coding novelty in Drosophila. Journal of molecular evolution 88, 382–398 (2020).

57. Lange, A. et al. Structural and functional characterization of a putative de novo gene in Drosophila. Nature communications 12, 1–13 (2021).

58. Bornberg-Bauer, E., Hlouchova, K. & Lange, A. Structure and function of naturally evolved de novo proteins. Current Opinion in Structural Biology 68, 175–183 (2021).

59. Reinhardt, J. A. et al. De Novo ORFs in Drosophila Are Important to Organismal Fitness and Evolved Rapidly from Previously Non-coding Sequences. PLOS Genetics 9, e1003860 (2013).

60. Xie, C. et al. A de novo evolved gene in the house mouse regulates female pregnancy cycles. eLife 8, e44392 (2019).

61. Wang, M. et al. PaxDb, a database of protein abundance averages across all three domains of life. Molecular & cellular proteomics 11, 492–500 (2012).

62. Kozak, M. An analysis of 5’-noncoding sequences from 699 vertebrate messenger RNAs. Nucleic Acids Research 15, 8125–8148 (1987).

63. Kozak, M. Adherence to the first-AUG rule when a second AUG codon follows closely upon the first. Proceedings of the National Academy of Sciences 92, 7134 (1995).

64. Hinnebusch Alan G. Molecular Mechanism of Scanning and Start Codon Selection in Eukaryotes. Microbiology and Molecular Biology Reviews 75, 434–467 (2011).

65. Kearse, M. G. & Wilusz, J. E. Non-AUG translation: a new start for protein synthesis in eukaryotes. Genes & development 31, 1717–1731 (2017).

66. Cao, X. & Slavoff, S. A. Non-AUG start codons: Expanding and regulating the small and alternative ORFeome. Experimental Cell Research 391, 111973 (2020).

67. Clements J M, Laz T M, & Sherman F. Efficiency of translation initiation by non-AUG codons in Saccharomyces cerevisiae. Molecular and Cellular Biology 8, 4533–4536 (1988).

68. Ivanov, I. P., Firth, A. E., Michel, A. M., Atkins, J. F. & Baranov, P. V. Identification of evolutionarily conserved non-AUG-initiated N-terminal extensions in human coding sequences. Nucleic Acids Research 39, 4220–4234 (2011).

69. Barahimipour, R. et al. Dissecting the contributions of GC content and codon usage to gene expression in the model alga Chlamydomonas reinhardtii. The Plant Journal 84, 704–717 (2015).

70. Belinky, F., Rogozin, I. B. & Koonin, E. V. Selection on start codons in prokaryotes and potential compensatory nucleotide substitutions. Scientific Reports 7, 12422 (2017).

71. Radío, S., Garat, B., Sotelo-Silveira, J. & Smircich, P. Upstream ORFs Influence Translation Efficiency in the Parasite Trypanosoma cruzi. Frontiers in Genetics 11, (2020).

72. Andreev, D. E. et al. Non-AUG translation initiation in mammals. Genome Biology 23, 111 (2022).

73. Gardner, L. B. Hypoxic inhibition of nonsense-mediated RNA decay regulates gene expression and the integrated stress response. Molecular and cellular biology 28, 3729– 3741 (2008).

74. Zetoune, A. B. et al. Comparison of nonsense-mediated mRNA decay efficiency in various murine tissues. BMC genetics 9, 1–11 (2008).

75. Heinen, T. J., Staubach, F., Häming, D. & Tautz, D. Emergence of a new gene from an intergenic region. Current Biology 19, 1527–1531 (2009).

76. Xie, C. et al. Hominoid-specific de novo protein-coding genes originating from long non-coding RNAs. PLoS genetics 8, (2012).

77. Papadopoulos, C., Chevrollier, N. & Lopes, A. Exploring the Peptide Potential of Genomes. in Computational Peptide Science. Methods in Molecular Biology (ed. Simonson, T.) vol. 2405 63–82 (Springer US, 2022).

78. Cherry, J. M. et al. Saccharomyces Genome Database: the genomics resource of budding yeast. Nucleic acids research 40, D700–D705 (2012).

79. Leinonen, R., Sugawara, H., Shumway, M., & on behalf of the International Nucleotide Sequence Database Collaboration. The Sequence Read Archive. Nucleic Acids Research 39, D19–D21 (2011).

80. Martin, M. Cutadapt removes adapter sequences from high-throughput sequencing reads. EMBnet. journal 17, 10–12 (2011).

81. Kim, D., Paggi, J. M., Park, C., Bennett, C. & Salzberg, S. L. Graph-based genome alignment and genotyping with HISAT2 and HISAT-genotype. Nature Biotechnology 37, 907–915 (2019).

82. Langmead, B. & Salzberg, S. L. Fast gapped-read alignment with Bowtie 2. Nature Methods 9, 357–359 (2012).

83. Lauria, F. et al. riboWaltz: Optimization of ribosome P-site positioning in ribosome profiling data. PLOS Computational Biology 14, e1006169 (2018).

84. Faure, G. & Callebaut, I. Identification of hidden relationships from the coupling of hydrophobic cluster analysis and domain architecture information. Bioinformatics 29, 1726–1733 (2013).

85. Faure, G. & Callebaut, I. Comprehensive repertoire of foldable regions within whole genomes. PLoS computational biology 9, (2013).

86. Bitard-Feildel, T. & Callebaut, I. HCAtk and pyHCA: A Toolkit and Python API for the Hydrophobic Cluster Analysis of Protein Sequences. bioRxiv 249995 (2018).

87. Lamiable, A. et al. A topology-based investigation of protein interaction sites using Hydrophobic Cluster Analysis. Biochimie 167, 68–80 (2019).

88. Altschul, S. F., Gish, W., Miller, W., Myers, E. W. & Lipman, D. J. Basic local alignment search tool. Journal of molecular biology 215, 403–410 (1990).

89. Hedges, S. B., Dudley, J. & Kumar, S. TimeTree: a public knowledge-base of divergence times among organisms. Bioinformatics 22, 2971–2972 (2006).

90. Kumar, S. et al. TimeTree 5: An Expanded Resource for Species Divergence Times. Molecular Biology and Evolution 39, msac174 (2022).

91. Team R Core, R. C. R: A language and environment for statistical computing. (2020).

92. Lin, M., Lucas Jr, H. C. & Shmueli, G. Research commentary—too big to fail: large samples and the p-value problem. Information Systems Research 24, 906–917 (2013).

93. Szabó, Á. et al. Ubiquitylation Dynamics of the Clock Cell Proteome and TIMELESS during a Circadian Cycle. Cell Reports 23, 2273–2282 (2018).

94. Perkins, D., Pappin, D., Creasy, D. & Cottrell, J. Probability-based protein identification by searching sequence databases using mass spectrometry data. Electrophoresis 20, 3551–3567 (1999).

95. Elias, J. E. & Gygi, S. P. Target-decoy search strategy for increased confidence in large-scale protein identifications by mass spectrometry. Nature Methods 4, 207–214 (2007).

96. Benjamini, Y. & Hochberg, Y. Controlling the False Discovery Rate: A Practical and Powerful Approach to Multiple Testing. Journal of the Royal Statistical Society. Series B (Methodological) 57, 289–300 (1995).

## References

Altschul SF, Gish W, Miller W, Myers EW, Lipman DJ. 1990. Basic local alignment search tool. Journal of molecular biology 215: 403–410.

Cherry JM, Hong EL, Amundsen C, Balakrishnan R, Binkley G, Chan ET, Christie KR, Costanzo MC, Dwight SS, Engel SR. 2012. Saccharomyces Genome Database: the genomics resource of budding yeast. Nucleic acids research 40: D700–D705.

Hedges SB, Dudley J, Kumar S. 2006. TimeTree: a public knowledge-base of divergence times among organisms. Bioinformatics 22: 2971–2972.

Kim D, Paggi JM, Park C, Bennett C, Salzberg SL. 2019. Graph-based genome alignment and genotyping with HISAT2 and HISAT-genotype. Nature Biotechnology 37: 907–915.

Kumar S, Suleski M, Craig JM, Kasprowicz AE, Sanderford M, Li M, Stecher G, Hedges SB. 2022. TimeTree 5: An Expanded Resource for Species Divergence Times. Molecular Biology and Evolution 39: msac174.

Langmead B, Salzberg SL. 2012. Fast gapped-read alignment with Bowtie 2. Nature Methods 9: 357–359.

Lauria F, Tebaldi T, Bernabò P, Groen EJN, Gillingwater TH, Viero G. 2018. riboWaltz: Optimization of ribosome P-site positioning in ribosome profiling data. PLOS Computational Biology 14: e1006169.

Martin M. 2011. Cutadapt removes adapter sequences from high-throughput sequencing reads. EMBnet journal 17: 10–12.

Papadopoulos C, Chevrollier N, Lopes A. 2022. Exploring the peptide potential of genomes. In Computational Peptide Science, Methods in Molecular Biology (ed. T. Simonson), Vol. 2405 of, Springer US.

